# PKR drives neuropathology in a mouse model of Leigh syndrome

**DOI:** 10.64898/2026.11.10.765677

**Authors:** Melania González-Torres, Patrizia Bianchi, Marcos Blanco-Ramos, Marta Luna-Sánchez, Patricia Prada-Dacasa, Emma Puighermanal, Carla Canal, Laura Cutando, Stephanie Le, Álex Gella, Mònica Girona, Laura Sánchez-Benito, Berta Armengol, Isabella Appiah, Joaquín Fernández-Irigoyen, Mohammed Alkhatib, Leonard Feist, Gregory McElroy, Pawel Lisowski, Antonio del Sol, Enrique Santamaría, Alessandro Prigione, Elisenda Sanz, Albert Quintana

## Abstract

Mitochondrial dysfunction is increasingly recognized as a trigger of innate immune responses, yet whether these responses contribute to neurodegeneration remains poorly understood. Here, we uncover a cell-type-specific antiviral program in vulnerable GABAergic neurons of *Ndufs4*-deficient mice and human brain organoids modeling Leigh syndrome. Despite robust type I interferon responses, genetic ablation of innate immune mediators *Ifnar1* or *Sting1* fails to ameliorate disease progression. Instead, we identify the double-stranded RNA sensor protein kinase R (PKR) as a critical mediator of mitochondrial neuropathology. PKR associates with mitochondrial RNA, while its genetic deletion or silencing delays neurological decline, attenuates neuroinflammation, and extends survival without restoring mitochondrial respiration. PKR deletion also normalizes JNK signaling without detectable global translational repression, suggesting a noncanonical contribution to disease. Together, our findings reveal PKR-mediated antiviral RNA sensing as a pathogenic mechanism in mitochondrial complex I deficiency and highlight PKR as a candidate target for mitochondrial encephalopathies.

## Introduction

Neurons are highly dependent on mitochondrial function to meet their exceptional energetic and metabolic demands, making them particularly vulnerable to mitochondrial dysfunction (Camandola and Mattson, 2017; Lax et al., 2017). This vulnerability is best illustrated by primary mitochondrial diseases (MDs), a group of largely untreatable genetic disorders that affect the mitochondrial energy-generating machinery and frequently lead to neurological dysfunction (Gorman et al., 2016; Gorman et al., 2015; McFarland et al., 2010; Rahman and Rahman, 2018). Among them, Leigh syndrome (LS), the most common pediatric MD, is characterized by progressive neurodegeneration, focal brain lesions, and early mortality (Lake et al., 2016; Magro et al., 2025). Despite being caused by mutations in more than 100 genes affecting mitochondrial function, LS shows striking anatomical and cell-type specificity. However, the mechanisms underlying this selective vulnerability remain poorly understood (Bolea et al., 2019; Fassone and Rahman, 2012; Lake *et al*., 2016; Quintana et al., 2010; Sanchez-Benito et al., 2025).

Beyond bioenergetic failure, MD pathogenesis involves oxidative stress, defective autophagy, and activation of cellular stress responses, among other mechanisms (Suomalainen and Battersby, 2018; van de Wal et al., 2022). More recently, immune activation has emerged as an important contributor to disease pathology (Aguilar et al., 2022; Hanaford and Johnson, 2022; Hanaford et al., 2023; Stokes et al., 2022). In this context, mitochondria have been identified as central regulators of innate immunity (Marques et al., 2024). Consistent with their endosymbiotic origin, mitochondria preserve bacterial traits and their components can be sensed by innate immune sensors, thereby initiating inflammatory and antiviral responses (Luna et al., 2025; Marques *et al*., 2024). Among these, type I interferon (IFN-I) signaling has received particular attention in MDs, where it has been proposed both as a biomarker and as a contributor to disease pathogenesis, primarily in peripheral tissues and in disease models lacking central nervous system involvement (Dhir et al., 2018; Keshavan et al., 2024; Lei et al., 2021; Lepelley et al., 2021; Warren et al., 2023). Recent studies have also identified IFN-I-related genes and proteins in samples from LS patients (Keshavan *et al*., 2024) and mouse models (Fouche et al., 2026; McElroy et al., 2020; Perry et al., 2021). However, the cellular sources and targets of this response, and its contribution to LS neuropathology, remain unknown. Establishing whether antiviral signaling originates within vulnerable neurons and contributes to disease progression is key to defining its role in LS neuropathology.

Here, we show that mitochondrial complex I (CI) deficiency induces an antiviral response in vulnerable neuronal populations in a mouse model of LS and human striatal organoids. We identify the double-stranded RNA sensor protein kinase R (PKR) as a key mediator of disease progression, revealing a previously unrecognized link between mitochondrial dysfunction, antiviral signaling and LS neuropathology.

## Results

### Ndufs4 deficiency induces an antiviral program in vulnerable brain areas

To investigate the basis of selective vulnerability in LS, we used the *Ndufs4* knockout (Ndufs4KO) mouse model, in which loss of the CI subunit *Ndufs4* recapitulates key disease features (Kruse et al., 2008; Quintana *et al*., 2010; van de Wal *et al*., 2022). Pathology in these mice emerges in a highly region-specific manner, showing gliosis, spongiform degeneration and neuronal loss (Johnson et al., 2013; Liu et al., 2015; Quintana *et al*., 2010). Within affected regions, GABAergic neurons in the olfactory bulb (OB) and globus pallidus externus (GPe) are selectively lost (Bolea *et al*., 2019; Sanchez-Benito *et al*., 2025), although the mechanisms underlying this vulnerability remain unresolved.

Quantitative proteomic analysis of OB samples from Ndufs4KO mice revealed a robust upregulation of proteins associated with IFN-I responses, including the interferon-stimulated gene (ISG) products ISG15 and IFIT3 (Figure 1A). Western blotting confirmed accumulation of both free and conjugated ISG15 proteins (ISGylation), a ubiquitin-like post-translational modification induced downstream of IFN-I signaling (Figure 1B), together with increased IFN-α levels measured by ELISA (Figure 1C), supporting activation of IFN-I signaling from early disease stages.

**Figure 1.**
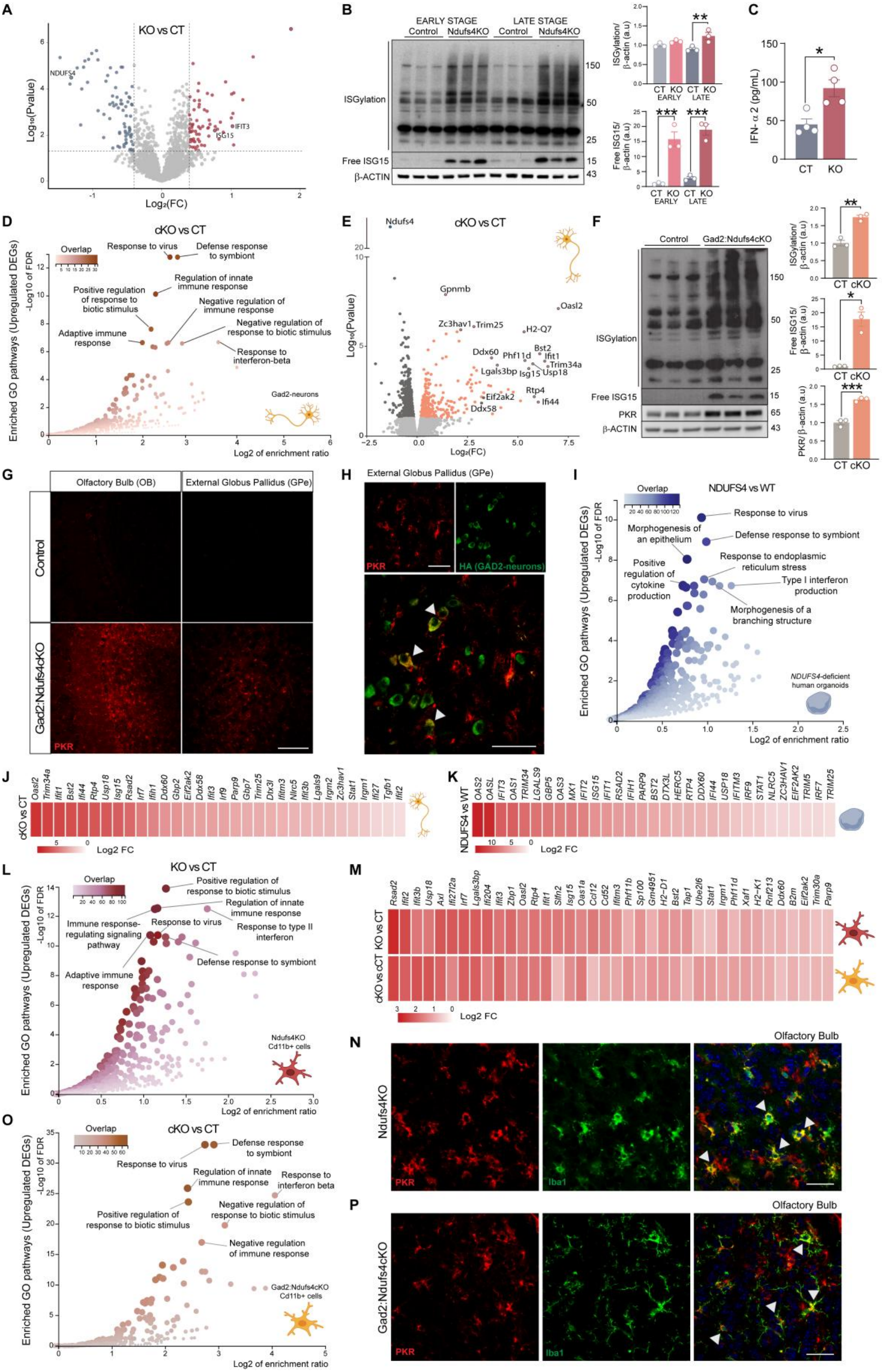
Antiviral-like signatures in Ndufs4KO brains and *NDUFS4*-deficient human striatal organoids. (A) Volcano plot of quantitative proteomics in the OB of Ndufs4KO (KO) vs control (CT) mice at late disease stage (n=4). (B) Representative immunoblot of ISGylated proteins and free ISG15 in the OB of control and Ndufs4KO mice at early and late disease stages, with quantification of ISGylation and free ISG15 normalized to β-ACTIN (n=3). Mean ± SEM; two-way ANOVA followed by Tukey’s post hoc. **p < 0.01, ***p < 0.001. (C) IFN-α2 levels in the OB measured by ELISA (n=4). Mean ± SEM; unpaired Student’s t-test. *p < 0.05. (D) Gene Ontology (biological process) enrichment of upregulated DEGs in the ribosome-associated transcriptome of OB GABAergic neurons from Gad2:RiboTag:Ndufs4cKO vs control mice (cKO vs CT; n=3). (E) Volcano plot of the same comparison highlighting antiviral and IFN-I-related transcripts. (F) Representative immunoblot and quantification of ISGylation, free ISG15 and PKR in the OB of Gad2:Ndufs4cKO mice, normalized to β-ACTIN (n=3). Mean ± SEM; unpaired Student’s t-test. *p < 0.05, **p < 0.01, ***p < 0.001. (G) Representative confocal images of PKR immunostaining (red) in the OB and globus pallidus externus (GPe) of control and Gad2:RiboTag:Ndufs4cKO mice. Scale bar, 150 µm. (H) Higher-magnification images of PKR (red) and HA (green; *Gad2*-expressing neurons) in the GPe of Gad2:RiboTag:Ndufs4cKO mice; arrowheads indicate representative double-positive neurons. Scale bars, 30 µm. (I) GO (biological process) enrichment of upregulated genes in *NDUFS4*-deficient human striatal organoids (NDUFS4 vs WT, n=2). (J) Heatmap (Log_2_FC) of antiviral genes in *Ndufs4*-deficient OB GABAergic neurons (cKO vs CT). (K) Heatmap (Log_2_FC) of antiviral genes in *NDUFS4*-deficient human striatal organoids (NDUFS4 vs WT, n=2). (L) GO enrichment of upregulated genes in CD11b⁺ cells from the OB of Ndufs4KO mice (KO vs CT; n=5 biological replicates; each pooled from 3-4 animals). (M) Heatmap (Log_2_FC) of IRM signature genes in CD11b⁺ cells from Ndufs4KO (KO vs CT, upper row) and Gad2:Ndufs4cKO (cKO vs CT, lower row) mice. (N) PKR (red), IBA1 (green) and DAPI (blue) in the OB of Ndufs4KO mice; arrowheads indicate representative double-positive cells. Scale bar, 50 µm. (O) GO enrichment of upregulated genes in CD11b⁺ cells from the OB of Gad2:Ndufs4cKO mice (cKO vs CT; n=4-5 biological replicates; each pooled from 3-4 animals). (P) PKR (red), IBA1 (green) and DAPI (blue) in the OB of Gad2:Ndufs4cKO mice; arrowheads indicate representative double-positive cells. Scale bar, 30 µm.

To determine whether this antiviral response arises from vulnerable neuronal populations, we next profiled the cell type-specific translatome of GABAergic neurons from the OB using the RiboTag approach (Sanz et al., 2009) in mice with conditional *Ndufs4* deletion in *Gad2*-positive cells (Gad2:RiboTag:Ndufs4cKO) (Figure S1A-C). RNA-seq of HA-tagged ribosome-associated mRNAs (Figure S1D,E) revealed upregulation of canonical ISGs such as *Ifit1*, *Isg15*, and *Oasl2*, together with antiviral nucleic acid sensors such as *Eif2ak2* (encoding PKR), and *Ddx58* (encoding retinoic acid-inducible gene I, RIG-I) (Figure 1D,E). Gene ontology analysis of upregulated genes highlighted enrichment of “response to virus”, and “response to interferon-beta” categories (Figure 1D), indicating activation of an antiviral program in CI-deficient GABAergic neurons. Notably, this response was not shared by vestibular glutamatergic *Slc17a6*-expressing neurons, another vulnerable population previously characterized (Bolea *et al*., 2019). RiboTag profiling of vestibular glutamatergic neurons carrying the same conditional deletion (Vglut2:RiboTag:Ndufs4cKO), showed no correlation with the transcriptional response of GABAergic neurons (Spearman’s ρ = −0.033; Figure S1F), with antiviral genes induced in GABAergic neurons remaining unchanged in glutamatergic neurons. Thus, activation of an antiviral response appears to be cell type-specific rather than a generic consequence of *Ndufs4* deficiency.

These transcriptional changes observed in Gad2:Ribotag:Ndufs4cKO mice were accompanied by increased ISG15 and PKR protein levels (Figure 1F), and immunostaining revealed consistent PKR overexpression in GAD2-positive neurons from the GPe, another GABAergic region severely affected in Gad2:Ndufs4cKO mice (Figure 1G,H) (Bolea *et al*., 2019; Puighermanal et al., 2024; Sanchez-Benito *et al*., 2025).

To address whether these transcriptional responses to CI deficiency could be recapitulated in a human model, we differentiated induced pluripotent stem cells (iPSCs) carrying a knock-in *NDUFS4* pathogenic variant (c.316C>T) (Daneshgar et al., 2022; Zink et al., 2026) into striatal brain organoids (Miura et al., 2020), modelling a region commonly affected in LS individuals (Lake *et al*., 2016; Magro *et al*., 2025). Striatal organoids recapitulated the transcriptional antiviral response observed in the mouse model (Figure 1I,J,K), including marked upregulation of *ISG15, IFIT1, OASL* and *EIF2AK2*, demonstrating a conserved response across species.

We next asked whether antiviral signaling could be induced by acute *Ndufs4* loss, independently of the developmental and systemic consequences of constitutive CI deficiency. To that end, we acutely deleted *Ndufs4* in the adult OB via stereotaxic delivery of AAV1-CRE·GFP into *Ndufs4*-floxed animals (OB-cKO), targeting the granule cell layer, which is predominantly GABAergic (Figure S2A-D). Immunoprecipitation of ribosome-associated mRNAs from RiboTag-floxed mice injected with AAV1-CRE·GFP (Figure S2A) confirmed efficient recombination and depletion of *Ndufs4* mRNA in transduced cells (Figure S2B). Acute *Ndufs4* deletion increased antiviral protein levels at the injection site in OB-cKO mice compared with control mice injected with AAV1-ΔCRE·GFP, which expresses a non-functional Cre (Figure S2C-F). Thus, acute CI deficiency is sufficient to induce a local antiviral response in the adult brain.

Progressive microgliosis is a hallmark of LS (Daneshgar *et al*., 2022; Quintana *et al*., 2010), and microglia and peripheral myeloid cells have emerged as key contributors to Ndufs4KO pathogenesis (Aguilar *et al*., 2022; Hanaford and Johnson, 2022; Hanaford *et al*., 2023; Stokes *et al*., 2022). We therefore hypothesized that the antiviral program observed in GABAergic neurons in the Ndufs4KO mice could propagate to brain myeloid cells, driving them towards an interferon-responsive state. To that end, we profiled MACS-sorted CD11b⁺ cells from the OB of Ndufs4KO mice (Figure S3A-D), which showed enrichment of antiviral genes and a transcriptional profile consistent with interferon-responsive microglia (IRM) (Lopez-Atalaya and Bhojwani-Cabrera, 2025) (Figure 1L,M). Immunostaining of OB sections further revealed PKR expression in IBA1+ cells (Figure 1N). A similar signature was observed in CD11b⁺ cells from GABAergic neuron-specific Ndufs4cKO mice (Gad2:Ndufs4cKO) (Figure 1O,M) and was further supported by PKR immunostaining (Figure 1P). These results indicate that neuronal *Ndufs4* deficiency is sufficient to induce an IRM state in brain myeloid cells.

### IFN-I signaling shapes the immune response without affecting disease progression in Ndufs4KO mice

To determine the contribution of IFN-I signaling to disease pathogenesis, we generated *Ifnar1:Ndufs4* double-knockout (Ifnar1DKO) mice. Loss of IFN-I signaling did not affect disease progression, as survival, weight loss, motor deficits, and respiratory dysfunction were comparable to Ndufs4KO mice (Figure 2A, Figure S4A-F). Thermal-induced seizure assays showed no differences at postnatal day (PND) 42 (Figure S4G-I), while seizure duration, but not frequency or severity, was increased at PND55 (Figure 2B-D). Likewise, *Ifnar1* deletion failed to restore mitochondrial respiratory function in OBs, as both ATP-linked and maximal respiration remained significantly reduced in Ifnar1DKO mice compared with controls, with no significant improvement relative to Ndufs4KO mice (Figure S4J-L).

**Figure 2.**
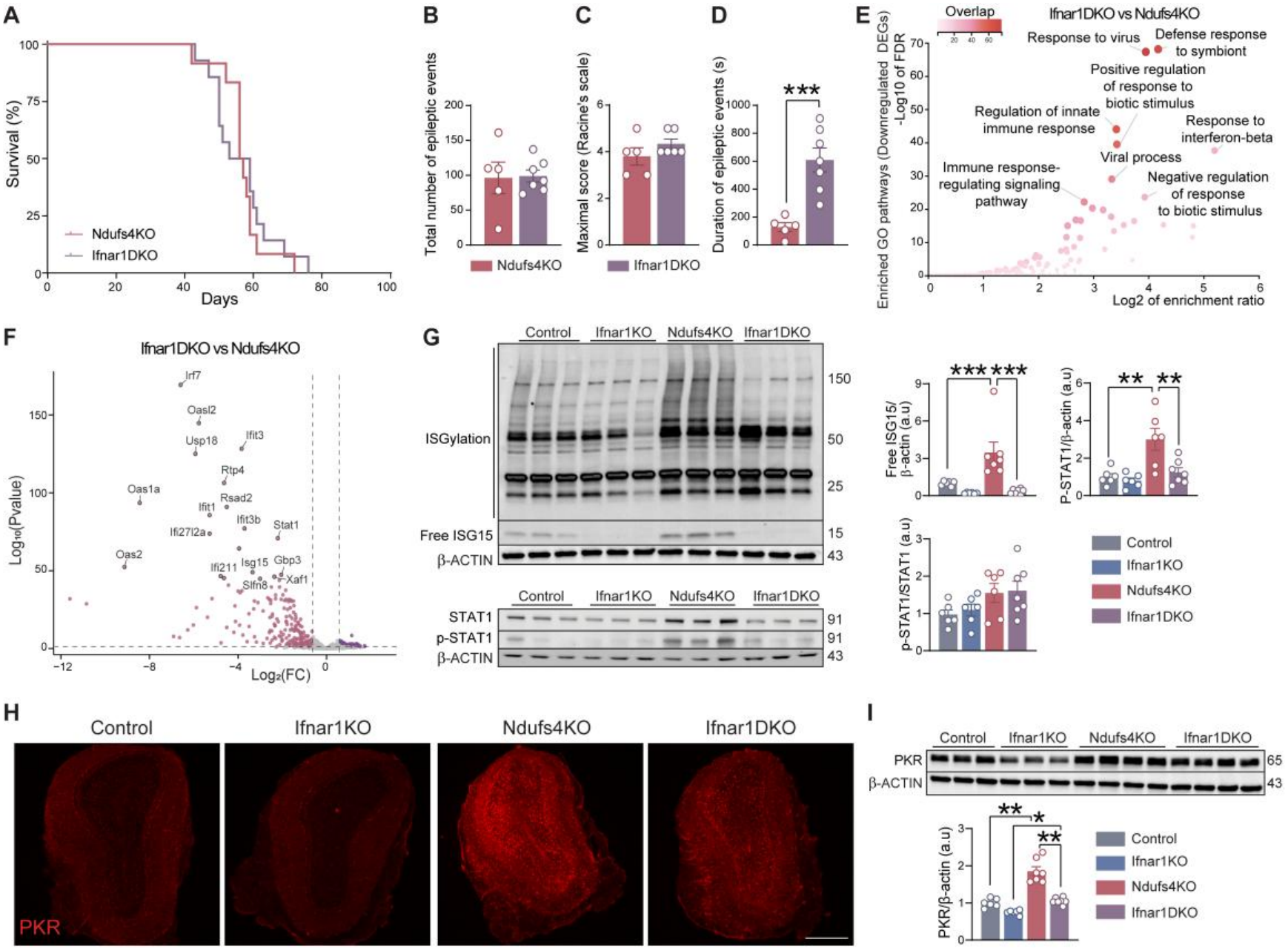
Blocking type I interferon signaling does not ameliorate the clinical phenotype of Ndufs4KO mice. (A) Survival curves of Ndufs4KO (n=12) and Ifnar1DKO mice (n=14). Log-rank test. (B-D) Heat-induced seizure susceptibility at late disease stage (PND55) in Ndufs4KO and Ifnar1DKO mice: (B) total number of epileptic events; (C) maximal seizure severity (modified Racine scale); (D) total duration of epileptic events (s). Mean ± SEM (n=6-7); unpaired Student’s t-test. ***p < 0.001. (E) GO enrichment of downregulated DEGs in the OB of Ifnar1DKO mice compared to Ndufs4KO mice (n=5 biological replicates; each pooled from 3-4 animals). (F) Volcano plot of the same comparison highlighting top downregulated genes. (G) Representative immunoblots of ISGylation/free ISG15 and of STAT1 and phospho-STAT1 (Tyr701) in the OB of all four genotypes. Quantification of free ISG15, p-STAT1 normalized to β-ACTIN, and p-STAT1/STAT1 ratio (n=6-7). Mean ± SEM; two-way ANOVA followed by Tukey’s post hoc. **p < 0.01, ***p < 0.001. (H) Representative immunofluorescence of PKR in the OB of control, Ifnar1KO, Ndufs4KO and Ifnar1DKO mice. Scale bar, 500 µm. (I) Representative immunoblot of PKR in all groups and quantification relative to β-ACTIN (n=6-7). Mean ± SEM; two-way ANOVA followed by Tukey’s post hoc. *p < 0.05, **p < 0.01.

At the molecular level, RNA-seq of OB samples (Figure S5A,B) showed that *Ifnar1* deletion markedly attenuated the canonical IFN-I transcriptional signature induced by *Ndufs4* loss (Figure 2E,F), including reduced expression of *Isg15, Ifit1* and *Oasl2* (Figure 2F) and decreased free ISG15 and phospho-STAT1 protein levels (Figure 2G). Despite this, genes belonging to immune and antimicrobial GO terms remained upregulated in Ifnar1DKO relative to Ifnar1KO mice (Figure S5C), as did antiviral nucleic acid sensors such as *Eif2ak2* (PKR) and *Sting1* (Figure S5D), albeit below Ndufs4KO levels. Total PKR protein was likewise increased in Ifnar1DKO versus Ifnar1KO mice (Figure 2H,I), indicating that *Ndufs4* deficiency induces PKR through both IFN-I-dependent and -independent mechanisms.

*Ifnar1* deletion also reduced microglial and macrophage markers (IBA1, CD11c, Cathepsin B) compared with Ndufs4KO mice (Figure 3A), with minimal changes in cytokine levels except for IL-6 (Figure S5E). RNA-seq of CD11b⁺ cells (Figure S5A,B) showed downregulation of ISGs and a near-complete loss of the Ndufs4KO IRM signature in Ifnar1DKO cells (Figure 3B,C), while comparison with Ifnar1KO cells confirmed that lack of IFNAR1 blunts the induction of IRM-associated genes and GO terms (Figure S6A). Accordingly, the induction of *Eif2ak2* observed in whole tissue samples was absent in Ifnar1DKO CD11b+ cells when comparing with Ifnar1KO group (Figure S6B,C). In line with this, PKR immunostaining revealed a reduction of PKR+ IBA1+ cells in Ifnar1DKO mice (Figure 3D), whereas PKR upregulation persisted predominantly in IBA1-negative cells (Figure S6D). Overall, IFN-I signaling shapes the microglial phenotype, whereas IFN-I-independent PKR induction occurs predominantly in non-microglial cells.

**Figure 3.**
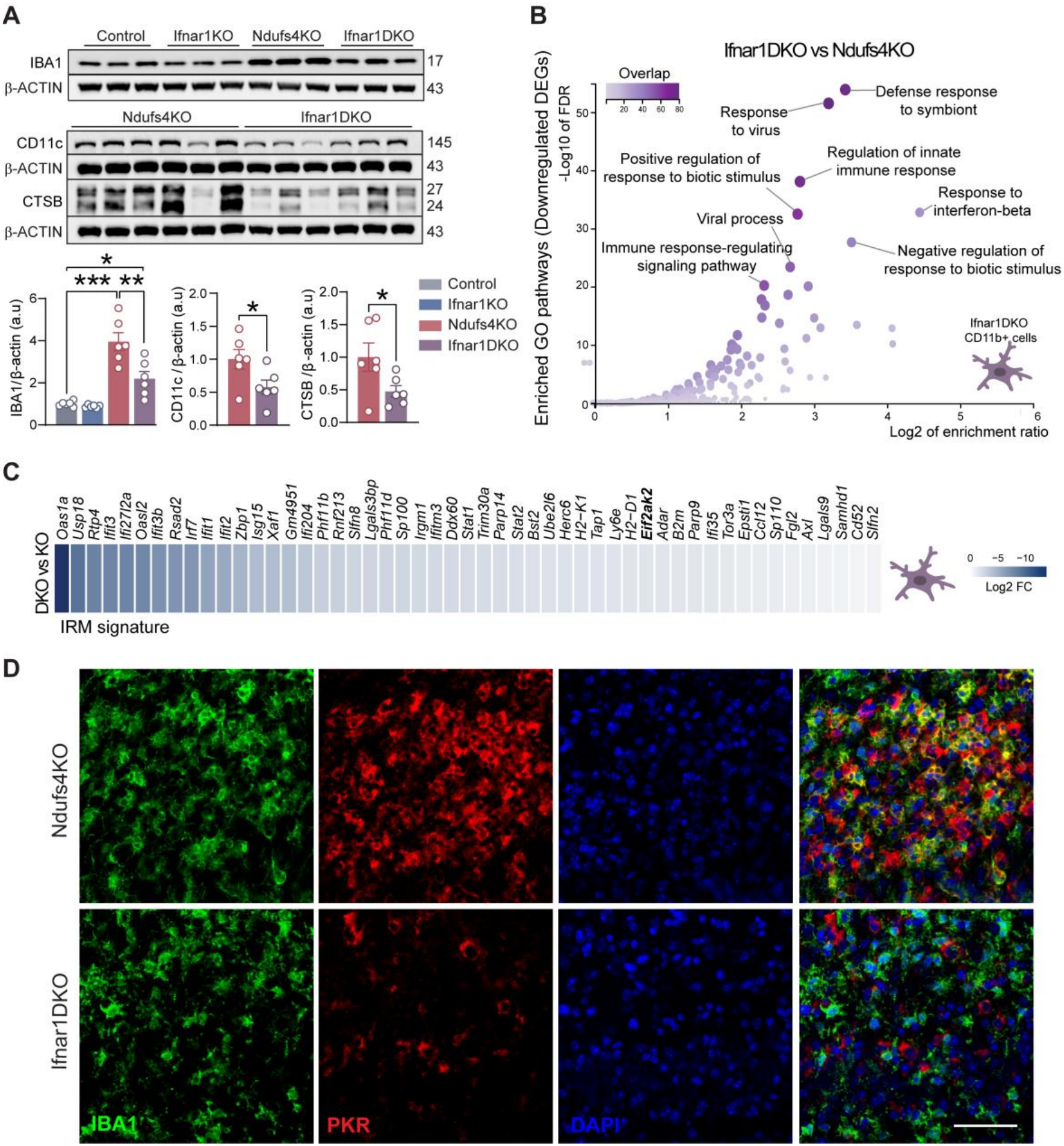
*Ifnar1* deletion prevents the IRM transcriptional signature. (A) Representative immunoblots of IBA1 (all four genotypes) and of CD11c and Cathepsin B (CTSB) (Ndufs4KO vs Ifnar1DKO) in the OB, with quantification normalized to β-ACTIN (n=6). Mean ± SEM; two-way ANOVA followed by Tukey’s post hoc for IBA1 and unpaired Student’s t-test for CD11c and CTSB. *p < 0.05, **p < 0.01, ***p < 0.001. (B) GO enrichment of downregulated genes in CD11b⁺ cells from the OB of Ifnar1DKO vs Ndufs4KO mice (n=5 biological replicates; each pooled from 3-4 animals). (C) Heatmap (Log_2_FC) of IRM signature genes downregulated in Ifnar1DKO vs Ndufs4KO CD11b⁺ cells. (D) IBA1 (green), PKR (red) and DAPI (blue) immunostaining in the OB of Ndufs4KO and Ifnar1DKO mice. Scale bars, 50 µm.

We therefore asked whether other nucleic acid-sensing pathways were induced by *Ndufs4* deficiency. STING (stimulator of interferon genes), a central mediator of cytosolic nucleic acid sensing and previously implicated in mitochondrial stress responses (West et al., 2015), was induced in Ndufs4KO mice (Figure S7A). However, genetic ablation of *Sting1* in Ndufs4KO mice failed to improve survival, with a modestly shortened lifespan in STING1DKO mice (Figure S7B). As with IFN-I signaling, STING activation accompanies *Ndufs4* deficiency but is not required for disease progression.

### PKR activation engages non-canonical signaling in Ndufs4KO mice

Based on these findings, we next focused on PKR as a potential mediator of Ndufs4KO neuropathology. Both total and phosphorylated PKR levels, the latter corresponding to the active form of the kinase, were robustly increased in the OB of Ndufs4KO mice (Figure 4A). PKR is activated by both viral double-stranded RNA (dsRNA) and endogenous mitochondrial dsRNA (mtdsRNA) (Dhir *et al*., 2018; Kim et al., 2018). To characterize RNA species associated with PKR *in vivo*, we immunoprecipitated PKR from Ndufs4KO brains and profiled PKR-bound RNA by dsRNA sequencing (Figure 4B,C; Figure S8A,B). This analysis identified a diverse set of nuclear and mitochondrial transcripts associated with PKR. Among these, *mt-Nd5* and *mt-Nd6* were significantly enriched in Ndufs4KO mice compared with controls (Figure 4C, S8C). Notably, mitochondrial reads mapped to the *mt-Nd5/mt-Nd6* locus (Figure 4B) (∼11.7-14.1 kb in the mouse mitochondrial genome), a region transcribed from both strands and previously shown to be a hotspot for dsRNA formation in human cells (Dhir *et al*., 2018; Kim *et al*., 2018). These findings indicate increased association of PKR with mitochondrial RNA in the context of CI deficiency.

**Figure 4.**
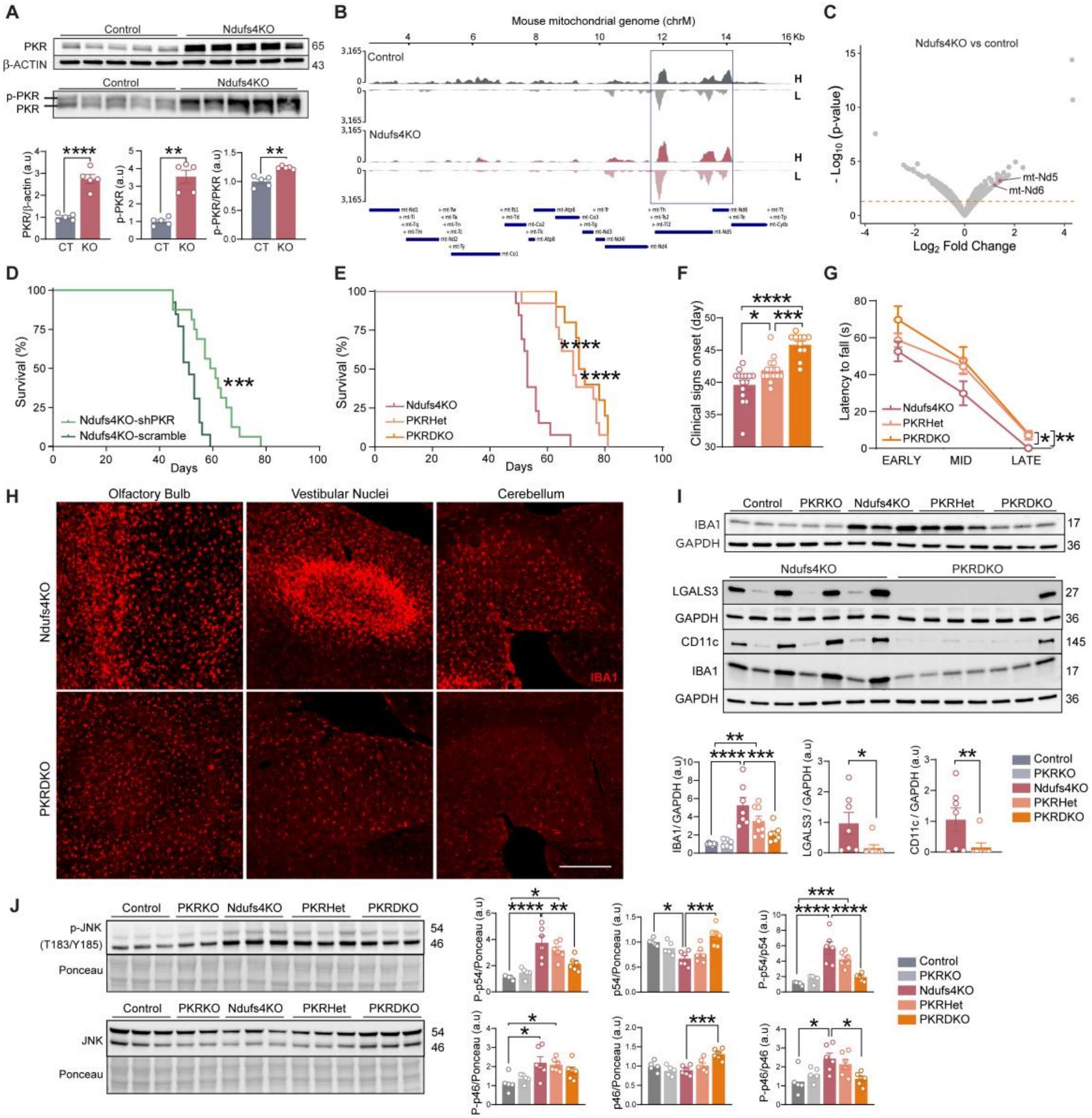
PKR drives neuroinflammation and disease progression in Ndufs4KO mice. (A) Representative immunoblots and quantification of total PKR (normalized to β-ACTIN), phosphorylated PKR (p-PKR, resolved by Phos-tag SDS-PAGE) and the p-PKR/PKR ratio in the OB of control and Ndufs4KO mice (n=5). Mean ± SEM; unpaired Student’s t-test. **p < 0.01, ****p < 0.0001. (B) Strand-specific coverage of PKR-bound RNA over the *mt-Nd1-mt-Nd6* interval of the mouse mitochondrial genome (GRCm39) in control and Ndufs4KO samples (H, heavy strand; L, light strand). The box indicates the 11.7-14.1 kb dsRNA hotspot (n=3). (C) Volcano plot of PKR-bound transcripts in Ndufs4KO vs control mice, with *mt-Nd5* and *mt-Nd6* highlighted. (D) Survival curves of scramble-injected (n=13) and AAV9-shPKR-injected (n=16) Ndufs4KO mice. Log-rank test. ***p < 0.001. (E) Survival curves of Ndufs4KO (n=13), PKRHet (n=13) and PKRDKO (n=10) mice. Log-rank test. ****p < 0.0001. (F) Onset of clinical signs in the same groups (n=12-15). Mean ± SEM; one-way ANOVA followed by Tukey’s post hoc. *p < 0.05, ***p < 0.001, ****p < 0.0001. (G) Latency to fall in the accelerating rotarod at early, mid and late disease stages (n=11-16). Mean ± SEM; mixed-effects model followed by Tukey’s post hoc. *p < 0.05, **p < 0.01. (H) Representative immunofluorescence of IBA1 in the OB, vestibular nuclei and cerebellum of Ndufs4KO and PKRDKO mice. Scale bar, 150 µm. (I) Representative immunoblots of IBA1 (all five genotypes) and of CD11c and Galectin-3 (LGALS3) (Ndufs4KO vs PKRDKO) in the OB normalized to GAPDH (n=7-8). Mean ± SEM; one-way ANOVA followed by Tukey’s post hoc for IBA1, Student’s t-test with Welch’ correction for LGALS3 and Mann-Whitney test for CD11c. *p < 0.05, **p < 0.01, ***p < 0.001, ****p < 0.0001. (J) Representative immunoblots of phospho-JNK (Thr183/Tyr185) and total JNK in the OB of all five genotypes, with quantification of phospho-JNK and total JNK normalized to Ponceau S, and of the p-p54/p54 and p-p46/p46 ratios (n=5-7). p54 and p46 denote the two JNK immunoreactive bands, quantified separately. Mean ± SEM; one-way ANOVA followed by Tukey’s post hoc. *p < 0.05, **p < 0.01, ***p < 0.001, ****p < 0.0001.

One of the best-characterized downstream functions of PKR is the inhibition of protein synthesis through phosphorylation of eIF2α at Ser51 (Gal-Ben-Ari et al., 2018). However, despite PKR activation, p-eIF2α levels were comparable between Ndufs4KO and control mice, as determined by both Phos-tag SDS-PAGE and immunoblotting with a phospho-specific antibody (Figure S9A). Consistently, bio-orthogonal non-canonical amino acid tagging (BONCAT) (Evans et al., 2020) revealed no evidence of global translational repression in Ndufs4KO mice (Figure S9B,C). Besides regulating translation, PKR can activate pro-inflammatory signaling cascades including JNK and NF-κB (Gal-Ben-Ari *et al*., 2018). Accordingly, both pathways appeared active in the OB of Ndufs4KO mice, with increased phosphorylation of the NF-κB subunit p65 (Ser536) and of JNK (Thr183/Tyr185) despite reduced total JNK protein levels (Figure S9D), as previously reported in this model (Liu *et al*., 2015).

### PKR is a causal mediator of neuropathology in Ndufs4KO mice

To determine the pathological relevance of PKR activation, we proceeded to silence PKR expression by means of a dual-route AAV9-shPKR-GFP delivery at PND1 (Corra et al., 2022). This approach resulted in widespread brain transduction (Figure S10A) and efficient PKR knockdown in both control and Ndufs4KO mice (Figure S10B). PKR silencing significantly extended survival and delayed disease onset in shPKR-injected compared to scramble-injected Ndufs4KO mice (Figure 4D, Figure S10C), with a modest improvement in motor performance (Figure S10D,E). Moreover, lower PKR protein levels were associated with longer survival in shPKR-treated mice, but not in scramble-treated controls (Figure S10F,G), linking PKR abundance to disease severity.

To assess whether complete genetic ablation of PKR recapitulated the protective effects of AAV-mediated knockdown, we generated a *Ndufs4:Eif2ak2* double-knockout mouse line. In agreement with our shRNA-mediated knockdown, genetic ablation of PKR significantly extended survival and delayed disease onset in PKRDKO (*Ndufs4*^Δ/Δ^:*Eif2ak2*^Δ/Δ^) mice. Notably, PKRHet (*Ndufs4*^Δ/Δ^:*Eif2ak2*^Δ/+^) mice showed a similar extension of survival but an intermediate delay in the onset of clinical signs, suggesting a gene-dosage effect on disease onset (Figure 4E,F). This phenotypic improvement in PKRDKO mice was accompanied by improved motor coordination (Figure 4G) and reduced body weight loss (Figure S11A,B). Consistent with an attenuated clinical phenotype, none of the PKRDKO mice were found dead, and only 10% of PKRDKO mice reached the predefined endpoint because of a severe phenotype, compared with 46% of Ndufs4KO mice that were either found dead or reached this endpoint. The remaining PKRDKO animals reached the predefined weight loss cut-off (Figure S11C). In contrast, PKR deletion did not affect locomotor activity in either Ndufs4KO (Figure S11D) or control mice (Figure S11E), nor did it alter rotarod performance in control animals (Figure S11F).

We next examined whether these phenotypic improvements were accompanied by reduced neuropathology. Both AAV-mediated knockdown and constitutive deletion of PKR markedly reduced microgliosis, as demonstrated by lower IBA1 protein levels and reduced immunoreactivity in vulnerable brain regions (Figure 4H,I; Figure S12A,B). PKR deletion also reduced GFAP while increasing GAD67 levels, consistent with preservation of the GABAergic phenotype in the OB (Figure S12C) and reduced disease associated microglia (DAM) markers CD11c and Galectin-3 (Figure 4I). Nevertheless, most PKRDKO mice eventually developed the characteristic vestibular nuclei lesions associated with fatal respiratory dysfunction (Bolea *et al*., 2019; Quintana et al., 2012), although the proportion of affected animals at matched ages was markedly lower than in Ndufs4KO littermates (Figure S12D,E), consistent with delayed disease progression. Of note, these benefits were not accompanied by an improvement in mitochondrial function, as OB mitochondria from PKRDKO mice displayed respiration rates similar to those of Ndufs4KO mice (Figure S12F), indicating that PKR acts downstream of, rather than restoring, the mitochondrial defect.

Given that PKR can activate JNK, a stress-signaling pathway previously shown to be altered in Ndufs4KO mice (Liu *et al*., 2015), as well as other inflammatory pathways including NF-κB (Gal-Ben-Ari *et al*., 2018), we next examined whether these responses were affected by genetic PKR deletion. Absence of PKR normalized JNK signaling, restoring the p-JNK(Thr183/Tyr185)/JNK ratio to control levels and reversing the reduction in total JNK (Figure 4J). PKRHet mice showed intermediate values, paralleling their moderate phenotypic improvement. In contrast, the effects on NF-κB were less pronounced. PKR deletion showed a tendency towards reduction of total p65 observed in Ndufs4KO mice, whereas phosphorylated p65 (Ser536) levels and the p-p65/p65 ratio remained unchanged (Figure S12G). Thus, normalization of JNK signaling more closely paralleled the phenotypic improvement associated with PKR deletion. Together, our results establish PKR as a key mediator of Ndufs4KO pathogenesis and identify JNK signaling as a potential downstream mechanism, without evidence of canonical eIF2α-dependent translational repression.

## Discussion

Mitochondrial dysfunction is increasingly recognized as a driver of innate immune activation, but whether antiviral pathways actively drive mitochondrial encephalopathies rather than merely accompany disease progression remains poorly understood. Here we show that complex I deficiency engages a cell-type-specific antiviral response in vulnerable neuronal populations, and we identify the dsRNA sensor PKR as a causal mediator of neuropathology in the Ndufs4KO mouse model of LS. Silencing or deleting PKR extended survival, delayed disease onset, improved motor performance, and reduced microgliosis, with PKR haploinsufficiency conferring intermediate protection. To our knowledge, these findings provide the first evidence that an antiviral RNA sensor actively contributes to disease progression in a mitochondrial encephalopathy.

Despite a robust and widespread ISG signature, ablation of *Ifnar1* in Ndufs4KO mice did not impact survival, weight loss, motor deficits or respiratory dysfunction. Similarly, loss of STING failed to confer protection, and modestly shortened survival. In contrast, pharmacological STING inhibition has been recently reported to protect neuro-retinal structure and function in Ndufs4KO mice (Lee et al., 2026), suggesting that STING may contribute to neuropathology in a region-specific manner. Our findings also contrast with POLG-mutator mice and several neurodegenerative models in which IFNAR1 or STING loss is protective (Lei *et al*., 2021; Nazmi et al., 2019; Roy et al., 2022; Roy et al., 2020; Wang et al., 2011). Moreover, *Ifnar1* deletion modestly worsened the seizure phenotype, consistent with emerging roles for IFN-I in synaptic plasticity and excitability (Di Filippo et al., 2016; Hosseini et al., 2020). Together, these observations indicate that the prominent IFN-I response is not a major determinant of disease progression in this model and illustrate that the magnitude of an antiviral signature does not necessarily reflect its pathogenic relevance. PKR, in contrast, remained elevated in *Ifnar1*-deficient animals. Given its ability to respond to diverse cellular stress signals beyond interferons (Kang and Tang, 2012), PKR may provide a route through which mitochondrial dysfunction engages pathogenic signaling independently of STING and canonical IFN-I signaling. This mechanism differs from previously described mechanisms of innate immune activation in MD involving mitochondrial DNA release and cGAS-STING signaling (Lei *et al*., 2021; West *et al*., 2015).

Mitochondrial RNA has also been shown to elicit antiviral responses, particularly in settings involving direct defects in mitochondrial RNA processing or degradation (Dhir *et al*., 2018). Our findings extend this framework to CI deficiency, in which increased association of PKR with mtRNA occurs without a primary genetic defect in mitochondrial RNA-processing machinery. The enrichment of mt-*Nd5* and mt-*Nd6* among PKR-associated transcripts, together with reads mapping to the *mt-Nd5*/*mt-Nd6* locus, a described hotspot for dsRNA formation (Dhir *et al*., 2018; Kim *et al*., 2018), provides a potential molecular link between CI deficiency and endogenous RNA sensing. These observations support investigating mitochondrial dsRNA-PKR signaling as an alternative contributor to MD progression.

Selective neuronal vulnerability is a defining feature of LS, and *Ndufs4* deficiency selectively affects GABAergic neuronal populations in the OB and GPe (Bolea *et al*., 2019; Quintana *et al*., 2010; Sanchez-Benito *et al*., 2025; van de Wal *et al*., 2022). The differential antiviral responses of vulnerable GABAergic and glutamatergic neurons indicate that CI alone is insufficient to explain engagement of this transcriptional program. Instead, neuronal identity appears to influence how mitochondrial dysfunction is translated into antiviral signaling. PKR induction has also been described in affected neuronal populations in other neurodegenerative disorders (Bando et al., 2005; Bose et al., 2011; Onuki et al., 2004). In particular, the relationship between mitochondrial dysfunction, PKR and vulnerable GABAergic neurons in Huntington’s disease (Lee et al., 2020) may point to a broader connection between neuronal identity and antiviral stress responses. However, the cellular determinants underlying these divergent responses to mitochondrial dysfunction remain to be defined.

Microglia and peripheral myeloid cells are established contributors to Ndufs4KO pathogenesis (Aguilar *et al*., 2022; Hanaford *et al*., 2023; Stokes *et al*., 2022). Our findings suggest that neuronal mitochondrial dysfunction can propagate an antiviral response to the myeloid compartment, consistent with the emergence of an IRM-like transcriptional state following neuron-restricted *Ndufs4* deletion. Similar IRM states have been described in aging and neurodegeneration (Lopez-Atalaya and Bhojwani-Cabrera, 2025). However, the near-complete suppression of this state by *Ifnar1* deletion without a corresponding improvement in disease progression dissociates the IFN-I-dependent myeloid signature from the pathological outcomes examined. Together with previous evidence implicating microglia and peripheral macrophages in this model (Aguilar *et al*., 2022; Hanaford *et al*., 2023; Stokes *et al*., 2022), these observations suggest that additional myeloid responses, beyond the IFN-I-dependent IRM state, contribute to disease progression.

Mechanistically, the absence of detectable eIF2α phosphorylation or global translational repression suggests that PKR contributes to Ndufs4KO pathology through mechanisms beyond its canonical role in translational repression. Instead, PKR deletion normalized JNK signaling, reversing both the elevated p-JNK/JNK ratio and the reduction in total JNK observed in Ndufs4KO mice. These findings extend previous observations of JNK activation in Ndufs4KO mice and its contribution to mitochondrial dysfunction-associated neurodegeneration in *Drosophila* (Liu *et al*., 2015). In keeping with this, the attenuation of microgliosis and neuropathology following PKR knockdown or deletion suggests that PKR contributes to disease by activating inflammatory signaling rather than by canonical eIF2α phosphorylation, in line with findings from other neurodegenerative models (Tible et al., 2019). Together, these observations identify JNK as a candidate downstream pathway associated with PKR-mediated pathology, although its contribution to the protective effects of PKR deletion remains to be defined.

Collectively, our findings establish PKR as a functional link between CI deficiency and disease progression, distinguishing its pathogenic contribution from the prominent but non-beneficial IFN-I response. PKR inhibition has shown therapeutic benefits in other neurodegenerative models (Cheng et al., 2025; Lopez-Grancha et al., 2021; Tible *et al*., 2019). Thus, the protection conferred by both PKR knockdown and genetic deletion, including the intermediate benefit observed with haploinsufficiency, provides a rationale for exploring PKR-targeted strategies in mitochondrial encephalopathies.

## Methods

### Animals

All experimental procedures were performed in sex- and age-balanced groups of mice, with no sex differences observed. All procedures were approved by the Research Ethics Committee of the Universitat Autònoma de Barcelona (CERec) and authorized by Generalitat de Catalunya. Mice were group-housed under controlled environmental conditions, under a 12-h light-dark cycle at 22 ± 2°C and 55 ± 10% relative humidity, with *ad libitum* access to chow (Global Diet #2014S, Envigo) and water.

Experiments were performed on the following strains, where Δ denotes the constitutively deleted (null) allele: *Gad2*^Cre/+^ (GAD2-IRES-Cre, JAX #028867) (Taniguchi et al., 2011)), *Ifnar1*^Δ/Δ^ (JAX #010830) (Muller et al., 1994), RiboTag (*Rpl22*^HA/HA^, JAX #011029) (Sanz *et al*., 2009) and *Sting1^gt^* (JAX stock #017537; *goldenticket* loss-of-function allele) (Sauer et al., 2011) mice were obtained from The Jackson Laboratory. *Ndufs4*^lox/lox^ (JAX #026963) and *Ndufs4*^Δ/+^ (JAX #027058) lines were previously generated in our laboratory (Kruse *et al*., 2008; Quintana *et al*., 2010). *Slc17a6^iCre^* (BAC-Vglut2::Cre) (Borgius et al., 2010) mice were a gift from Dr Ole Kiehn (Karolinska Institutet, Sweden), and *Eif2ak2*^Δ/Δ^ animals (hereafter PKRKO; (Yang et al., 1995) from Dr Adolfo García-Sastre (Icahn School of Medicine at Mount Sinai, USA). Every strain was maintained on a pure C57BL/6J background through a minimum of ten backcross generations.

Constitutive knockout mice lacking *Ndufs4* (Ndufs4KO) were obtained as previously described (Kruse *et al*., 2008), with wild-type and heterozygous littermates as controls. Conditional knockouts lacking *Ndufs4* in GABAergic neurons (Gad2:Ndufs4cKO) were generated as described (Bolea *et al*., 2019), with *Gad2*^Cre/+^*Ndufs4*^lox/+^ littermates used as controls.

For cell-type-specific translatome profiling, these lines were crossed with RiboTag mice, which express a Cre-dependent HA-tagged ribosomal protein RPL22 (Sanz *et al*., 2009), to obtain *Gad2*^Cre/+:^*Rpl22*^HA/+^*Ndufs4*^lox/+^ (Gad2:RiboTag:Ndufs4cCT), and *Gad2^Cre^*^/+:^ *Rpl22^HA^*^/+^*Ndufs4^lox^*^/lox^ (Gad2:RiboTag:Ndufs4cKO). Similarly, *Slc17a6*^iCre^*Ndufs4*^Δ/+^ mice were crossed with *Ndufs4*^lox/+^:*Rpl22^HA^*^/HA^ mice to obtain Vglut2:RiboTag:Ndufs4cKO (*Slc17a6*^iCre^*Rpl22*^HA/+^*Ndufs4*^Δ/lox^) and Vglut2:RiboTag:Ndufs4cCT (*Slc17a6*^iCre^*Rpl22*^HA/+^*Ndufs4*^Δ/+^) mice. *Ndufs4*^lox/lox^:*Rpl22*^HA/HA^ mice and their *Ndufs4*^lox/+:^*Rpl22*^HA/HA^ controls were used for AAV-mediated *Ndufs4* deletion in the adult OB.

Double knockout lines were generated by crossing *Ifnar1*^Δ/Δ^, *Eif2ak2*^Δ/Δ^ or Sting1^gt^ with *Ndufs4*^Δ/+^ mice and intercrossing the resulting double heterozygotes. The *Ifnar1* cross yielded controls (*Ndufs4*^Δ/+^;*Ifnar1*^Δ/+^), Ifnar1KO (*Ndufs4*^+/+^:*Ifnar1*^Δ/Δ^ and *Ndufs4*^Δ/+^:*Ifnar1*^Δ/Δ^), Ndufs4KO (*Ndufs4*^Δ/Δ^;*Ifnar1*^Δ/+^) and Ifnar1DKO (*Ndufs4*^Δ/Δ^:*Ifnar1*^Δ/Δ^) mice. The PKR cross yielded control (*Ndufs4*^+/+^:*Eif2ak2*^+/+^), PKRKO (*Ndufs4*^+/+^:*Eif2ak2*^Δ/Δ^), Ndufs4KO (*Ndufs4*^Δ/Δ^:*Eif2ak2*^+/+^), PKRHet (*Ndufs4*^Δ/Δ^:*Eif2ak2*^Δ/+^) and PKRDKO (*Ndufs4*^Δ/Δ^:*Eif2ak2*^Δ/Δ^) mice. The Sting1 cross generated Ndufs4KO (*Ndufs4*^Δ/Δ^:*Sting1*^+/+^) and STING1DKO (*Ndufs4*^Δ/Δ^:*Sting1*^gt/gt^) mice.

Genotyping was performed by PCR using previously described primer sequences (Bolea *et al*., 2019; Kruse *et al*., 2008; Sanz *et al*., 2009). Ifnar1KO mice were genotyped using the primers from JAX Protocol 29816. *Eif2ak2* gene was genotyped by using the following sequences: Neo-P3 (ATTCGCAGCGCATCGCCTTCTATCGCC), PKR-C (ATGGCAGCCTACCTTCTGTCATG) and PKR-N (CTTCCTTTGCAGGCCACT GGGAG). Genotyping of *Sting1* was performed following JAX protocol 17625, based on Sanger sequencing of the PCR product generated with primers 25766 (forward, 5′-GAT CCG AAT GTT CAA TCA GC-3′) and 25767 (reverse, 5′-CGA TTC TTG ATG CCA GCA C-3′).

### Clinical monitoring and disease staging

Body weight and clinical status were recorded daily from weaning. Onset of clinical signs was defined as the first day on which clasping or abnormal twisting behaviors appeared (Quintana *et al*., 2010). In this work, early, mid and late disease stages correspond to PND29-31, PND38-41 and PND49-52, respectively. In survival experiments, mice were humanely euthanized according to the ethical committee protocol when body weight declined by more than 20% of maximum, or when presenting an endpoint phenotype (paw paralysis, severe ataxia or acute dehydration). Cause of death was classified as weight cut-off, endpoint phenotype or found dead. Euthanasia was performed by CO_2_ administration.

### Human iPSC-derived striatal organoids

Induced pluripotent stem cells (iPSCs) carrying a knock-in *NDUFS4* variant c.316C>T (NDUFS4_1; HVRDi004-B-6) and their isogenic wild-type (WT) control (PGP1; HVRDi004-B-7) were previously generated (Daneshgar *et al*., 2022). iPSCs were maintained feeder-free on Geltrex-coated plates (#A1569601, Thermo Fisher Scientific) in StemMACS iPS-Brew XF (#130-104-368, Miltenyi Biotec) supplemented with MycoZap Plus-CL (#VZA-2012, Lonza) at 37°C in a humidified atmosphere of 5% CO₂ and 5% O₂. Cells were passaged at 70-80% confluency with EDTA (0.5 mM, #15575-038, Invitrogen). ROCK inhibitor Y-27632 (10 µM, #ALX-270-333, Enzo Life Sciences) was added after passaging to promote survival. Cells were routinely tested for mycoplasma by PCR using nine primers detecting the six most common mycoplasma species.

Striatal organoids were generated following Miura et al. (2020), with modifications. iPSCs were dissociated into single cells with Accutase (#A6964, Sigma-Aldrich) for 7 min at 37°C, seeded at 3×10⁶ cells per well in AggreWell 800 24-well plates (#34815, STEMCELL Technologies) in iPS-Brew with 10 µM Y-27632, centrifuged at 100 × g for 3 min and incubated at 37°C with 5% CO₂. After 24 h, spheroids were harvested using a cut P1000 tip and transferred to 10 cm Petri dishes coated with Anti-Adherence Rinsing Solution (#7010, STEMCELL Technologies) containing neural induction medium (NIM), and maintained on an orbital shaker (#543-12310-00, Heidolph) at 37°C and 5% CO₂. NIM consisted of KnockOut DMEM (#10829-018, Gibco), 20% KnockOut Serum Replacement (#10828-028, Gibco), MEM non-essential amino acids (#11140-050, Gibco), sodium pyruvate (#11360-039, Gibco), GlutaMAX (1:100, #35050-061, Gibco) and penicillin-streptomycin (1:100, #15140-122, Gibco). During days 1-5, NIM was replaced daily and supplemented with dorsomorphin (2.5 µM, #P5499, Sigma-Aldrich) and SB-431542 (10 µM, #130-105-336, Miltenyi Biotec). From day 6, spheroids were cultured in neural medium (Neurobasal-A, #10888022; B-27 without vitamin A, #12587001; GlutaMAX and penicillin-streptomycin; all from Gibco) supplemented with IWR-1 (2.5 µM, #681669, Merck) and Activin A (50 ng/ml, #78001.1, STEMCELL Technologies). During days 11-21, SR11237 (100 nM, #3411, Tocris) was added to the same medium. From day 22 onward, organoids were maintained in neural medium supplemented with BDNF (20 ng/ml, #130-138-877, Miltenyi Biotec), NT-3 (20 ng/ml, #130-096-288, Miltenyi Biotec), sodium L-ascorbate (200 µM, #A4034, Sigma-Aldrich), dibutyryl-cAMP (db-cAMP; 100 µM, #100-0244, STEMCELL Technologies) and DHA (10 µM, #D2534, Sigma-Aldrich), with DAPT (2.5 µM, #130-110-489, Miltenyi Biotec) added from day 42. From day 46, organoids were maintained in Neurobasal-A with B-27 Plus (#17504001, Gibco), GlutaMAX and MycoZap Plus-CL. From day 6, medium was changed three times per week until day 46 and twice weekly thereafter.

### Viral vectors

pAAV-CRE-GFP and pAAV-ΔCRE-GFP plasmids were kindly provided by Prof. Richard Palmiter (University of Washington, USA) (Bruchas et al., 2011; Fadok et al., 2009). ΔCRE encodes a truncated, non-functional form of Cre used as a control. Recombinant AAV vectors encoding CRE or ΔCRE were generated in HEK293T cells with an AAV1 serotype, following established protocols (Quintana *et al*., 2012) and used at a titer of 2×10^9^ vg/µl. The AAV9-shPKR construct was designed through the VectorBuilder platform using an shRNA sequence targeting PKR (*Eif2ak2;* CCGGGGAGTAGCCATTACGTATAAACTCGAGTTTATACGTAATGGCTACTCC TTTTTG), kindly provided by Prof. Kobi Rosenblum (University of Haifa, Israel). AAV9-shPKR-GFP and AAV9-scramble-GFP preparations were produced by VectorBuilder at 10^13^ vg/ml.

### Neonatal AAV administration

Systemic gene silencing was performed by intravenous and intracerebroventricular injections of AAV9-shPKR-GFP or AAV9-scramble-GFP into PND1 pups, as described (Corra *et al*., 2022). Cryo-anaesthetized pups received 10^11^ vg in 40 μL into the left temporal vein, followed by 10^10^ vg in 4 μL into the right lateral ventricle, and were rewarmed before being returned to the dam.

### Stereotaxic surgery

To achieve AAV-mediated *Ndufs4* deletion in the adult OB, AAV1-CRE·GFP or AAV1-ΔCRE·GFP were injected bilaterally in *Ndufs4*-floxed mice at PND28-30. All surgical procedures were conducted under aseptic conditions. Mice were anesthetized with isoflurane (1.5-2% in O₂), and analgesia (ketoprofen, 5 mg/kg, subcutaneous; Sanofi-Aventis) and ocular protective gel (Viscotears, Bausch + Lomb) were administered before surgery. Body temperature and breathing were monitored throughout the procedure. Animals were placed in a robot-assisted stereotaxic frame (Neurostar, Germany), a small craniotomy was performed and AAVs were delivered according to Paxinos and Franklin (from Bregma in mm: AP +4.28 mm, ML ±0.6 mm, DV −1.8 mm and −2.8 mm; 0.5 µl per site at 0.1 µl/min) using a 32G blunt needle (#7803-04, Hamilton) coupled to a 5 µl Hamilton syringe (#7634-01). The needle was left in place for 5 min to allow diffusion and withdrawn at 1 mm/min. Animals were monitored until recovery and euthanized 60 days after surgery.

### Open Field

Locomotor activity was filmed with an overhead camera during a single 30-min session in a white acrylic arena (56 × 36.5 × 31 cm), at early, mid and late disease stages. Tracking and computation of total distance travelled were analyzed by using EthoVision XT 11.5 (Noldus) (Seibenhener and Wooten, 2015).

### Rotarod

Balance and motor coordination were assessed on an accelerating rod (LE8205, Panlab; 30 mm diameter) (Brooks and Dunnett, 2009) at the three disease stages. On day 1, mice were habituated to remain on the rod (30 mm diameter) for 60 s at 4 rpm. The test consisted of five trials spaced 7 min apart, with the rod accelerating from 4 to 40 rpm in steps of 2 rpm every 10 s. A trial finished when the animal dropped or reached the maximal time on the rod (5 minutes), and the five latencies were averaged.

### Whole-body plethysmography

Respiratory function was assessed by unrestrained whole-body plethysmography, using a 90-mm chamber (#PLY310, EMMS) (Prada-Dacasa et al., 2020). Each mouse explored the chamber for 45 min before a 15-min acquisition window, from which breathing rate (breaths/min) and tidal volume normalized to body weight (µl/g) were extracted.

### Thermal-induced seizures

Seizure threshold was determined at mid (PND42) and late (PND55) stages as reported (Sanchez-Benito *et al*., 2025; Stokes *et al*., 2022). After 10 min of acclimation at 37.5°C, core temperature was raised in 0.5°C steps every 2 min until a seizure occurred or 42°C was reached, with continuous readout from a rectal probe (#RET-4, PHYMEP) connected to a temperature controller (Physitemp Instruments). Animals were then cooled with a fan. Sessions were filmed and later scored blind for number of epileptic events, peak severity on a modified Racine (Sanchez-Benito *et al*., 2025) and cumulative event duration.

### Histology and immunofluorescence

Following euthanasia, brains were fixed overnight at 4°C in 4% (w/v) paraformaldehyde in phosphate-buffered saline (PBS). After cryoprotection in 30% (w/v) sucrose in PBS, brains were frozen on dry ice and coronal or sagittal sections (20-30 µm) were obtained using a cryostat (CM3050 S, Leica). Sections were permeabilized in PBS containing 0.2% (v/v) Triton X-100 (PBST), blocked for 1 h at room temperature (RT) in 10% (v/v) donkey serum in PBST, and incubated overnight at 4°C in 1% (v/v) donkey serum in PBST with the primary antibodies listed in Table S1. The following day, sections were washed in PBST and incubated for 1 h at RT with fluorophore-conjugated secondary antibodies (1:1,000; Table S1). After additional washes, sections were mounted and coverslipped with DAPI-containing mounting medium (Fluoromount-G, Electron Microscopy Sciences, #17984-24). Sections incubated with secondary antibodies only were included as negative controls in all experiments.

Fluorescence images were acquired on Nikon Eclipse 90i, EVOS M5000 or Zeiss LSM 700 microscopes. For PKR signal quantification, images were analyzed in ImageJ (National Institutes of Health, USA). PKR signal was quantified as integrated density within manually delimited regions of interest and normalized to area, averaging 2-7 sections per animal.

### Western blotting

OB tissue was homogenized in SDS sample buffer (62.5 mM Tris-HCl pH 6.8, 2% (w/v) SDS, 10% (v/v) glycerol). Protein concentration was measured with the Pierce BCA protein assay kit (#23225, Thermo Scientific). Thereafter, 15-20 µg of protein lysates were denatured with 50 mM DTT and 0.1% (w/v) bromophenol blue, subjected to 4-20% gradient SDS-PAGE (#4561094, Bio-Rad) and transferred to nitrocellulose membranes using the Trans-Blot Turbo RTA Mini Nitrocellulose Transfer Kit (#1704270, Bio-Rad). Membranes were blocked for 1 h with 5% (w/v) dried skimmed milk in Tris-buffered saline containing 0.1% Tween-20 (TBS-T) and incubated overnight at 4 °C with primary antibodies (Table S1). After incubation with the corresponding HRP-conjugated secondary antibodies (1:10,000; Jackson ImmunoResearch #111-035-003, #115-035-003 and #705-035-003), membranes were washed in TBS-T and developed using Clarity Western ECL Substrate (#1705060, Bio-Rad) on a ChemiDoc MP Imaging System (Bio-Rad) or by exposure to X-ray film. Bands were quantified using ImageJ software (National Institutes of Health, USA) or the Image Lab 6.1 Software (Bio-Rad) and normalized to β-ACTIN, GAPDH or total protein (Ponceau S staining, Sigma Aldrich, #P7170) as indicated in each figure legend.

### Phos-tag SDS-PAGE

Phosphorylation of PKR and eIF2α was additionally assessed by Phos-tag SDS-PAGE, which retards the migration of phosphorylated species and thus resolves them from their non-phosphorylated counterparts, allowing detection with antibodies against the total protein (Table S1). Samples were run on 12.5% Phos-tag SDS-PAGE gels (#195-17991, Wako) in Tris-glycine-SDS running buffer (25 mM Tris, 192 mM glycine, 0.1% (w/v) SDS) at 150 V for 3-5 h at 4°C. Gels were subsequently washed to chelate divalent cations, equilibrated, and proteins were transferred overnight by wet transfer according to the manufacturer’s instructions. Detection was performed as described for standard western blotting.

#### BONCAT

Newly synthesized proteins were analyzed by BONCAT following a protocol with modifications kindly provided by Dr. Liviu-Gabriel Bodea (University of Queensland, Australia) (Evans *et al*., 2020). Mice were fed a low-methionine diet (#TD110208, Envigo) for 7 days. On the day of the experiment, late-stage Ndufs4KO and control mice were injected intraperitoneally with 100 µg/g body weight of L-azidohomoalanine (AHA, Bio-Techne R&D Systems, 6584/10); AHA-naive littermates were processed in parallel as negative controls. Four hours later, OBs were collected and homogenized in 200 µl RIPA buffer supplemented with protease and phosphatase inhibitors (#04693132001, Roche). Protein was quantified by BCA assay and 50 µg per sample were conjugated to biotin alkyne using the Click-iT Protein Reaction Buffer Kit (#C10276, Invitrogen). Proteins were precipitated, washed, resuspended and resolved by SDS-PAGE. AHA-labelled proteins were detected with IRDye 800CW Streptavidin (#926-32230, LI-COR) and imaged using a LI-COR Odyssey Fc imaging system. Signals were quantified and normalized to total protein levels determined using REVERT 700 Total Protein Stain (LI-COR).

### Mitochondrial respiration analysis

Mitochondria were isolated from OBs by differential centrifugation as previously described (Gella et al., 2020). Isolated mitochondria (10–25 µg protein per well) were seeded into Seahorse XFp microplates in 25 µl of mitochondrial assay solution (MAS) supplemented with 0.5% (w/v) fatty acid-free bovine serum albumin (MAS-BSA) and immobilized by centrifugation at 2,000 × g for 20 min at 4 °C. Wells were subsequently supplemented with 155 µl of MAS-BSA containing 10 mM pyruvate and 5 mM malate. Oxygen consumption rate (OCR) was measured using a Seahorse XFp Extracellular Flux Analyzer (Agilent Technologies) under basal conditions and following sequential injections of ADP (5 mM), oligomycin (2 µM), FCCP (5 µM) and antimycin A/rotenone (2.5 µM and 2 µM, respectively). Three OCR measurements were recorded per condition and averaged for analysis. Basal respiration was calculated as baseline OCR minus antimycin A/rotenone-insensitive OCR; ATP-linked respiration as ADP-stimulated OCR minus oligomycin-insensitive OCR; and maximal respiration as FCCP-stimulated OCR minus antimycin A/rotenone-insensitive OCR.

### ELISA and multiplex cytokine assay

IFN-α levels were measured in OB homogenates using the Mouse Interferon alpha 2 ELISA Kit (#ab211648, Abcam). Pools of 5-6 OBs per group were homogenized in the cell extraction buffer provided with the kit, and supernatants adjusted to 7 µg/µl of protein were assayed following the manufacturer’s instructions.

Multiplex cytokine analysis was performed with a MILLIPLEX Mouse High Sensitivity T Cell Magnetic Bead Panel (#MHSTCMAG-70K, Millipore) to measure IL-6, TNF-α, IL-10, IFN-γ, CCL2, and CXCL1. Samples were homogenized, sonicated and centrifuged as described (Canal et al., 2026) and protein concentration was determined with the Pierce BCA Protein Assay Kit (#23225, Thermo Scientific). Data were acquired on a MAGPIX instrument (Luminex xMAP technology) and analyzed with xPONENT 4.2 software (Luminex Corporation). Cytokine levels were normalized to protein concentration; measurements falling below the first standard of the curve were recorded as 0, and undetectable analytes were excluded from plotting.

### Mass spectrometry based-quantitative proteomics

OB samples from late-stage Ndufs4KO and control mice (n=4 per group) were processed for iTRAQ-based quantitative proteomics as described (Unwin et al., 2010), with modifications. Briefly, proteins were extracted in urea/thiourea/CHAPS buffer, reduced with TCEP, alkylated with MMTS, digested with trypsin (1:20, w/w; Promega) and labelled with iTRAQ reagents (Sciex). The pooled peptides were fractionated by high-pH reversed-phase chromatography and analyzed by nano-LC-MS/MS on a 5600 TripleTOF mass spectrometer (AB Sciex). Data were processed with ProteinPilot 4.5 (Sciex) against the UniProt *Mus musculus* reference proteome (UP000000589), using only peptides unique to a given protein for quantification and a 1% global false discovery rate estimated by decoy database search. Proteins with an iTRAQ ratio < 0.77 or > 1.3 and P < 0.05 were considered differentially abundant.

### Tagged-ribosome immunoprecipitation and RNA extraction

HA-tagged-ribosome immunoprecipitation was performed as described (Sanz et al., 2019). Briefly, tissue was dissected from Gad2:RiboTag:Ndufs4cCT and Gad2:RiboTag:Ndufs4cKO mice (OB), Vglut2:RiboTag:Ndufs4cCT and Vglut2:RiboTag:Ndufs4cKO mice (vestibular nuclei), and from Ndufs4^lox/lox^:Rpl22^HA/HA^ and Ndufs4^lox/+^:Rpl22^HA/HA^ mice injected with AAV1-CRE·GFP, were homogenized by douncing in 1 ml of supplemented polysome buffer (Sanz *et al*., 2019). After centrifugation, 10% of the supernatant was saved as input fraction, while the remaining supernatant was incubated with 2.5 µg of anti-HA antibody (BioLegend, #901514) for 4 h at 4°C with constant gentle rotation, followed by overnight incubation at 4°C with 100 µl of protein A/G magnetic beads (#88803, Fisher Scientific). Immunoprecipitates were washed in high-salt buffer as described (Sanz *et al*., 2019), and total RNA was extracted from both input and immunoprecipitated (IP) fractions.

### CD11b^+^ cell isolation

OBs from 3-4 animals were pooled per sample and enzymatically dissociated using (Neural Tissue Dissociation Kit, Miltenyi Biotec #130-092-628) on a gentleMACS Octo Dissociator with heaters (Miltenyi Biotec #130-096-427), following the manufacturer’s instructions. Myelin was removed with Myelin Removal Beads II (Miltenyi Biotec #130-096-733) on LS columns (#130-042-401) and CD11b⁺ cells were positively selected with anti-CD11b magnetic microbeads (Miltenyi Biotec #130-093-636) on MS columns (#130-042-201). An aliquot of the dissociated tissue was retained before myelin removal as the whole-tissue fraction, and the flow-through was collected as the CD11b⁻ fraction.

### RNA extraction and quantification

Total RNA was extracted using the RNeasy Micro Kit with on-column DNase I treatment (#74104, Qiagen). RNA was quantified with the Quant-iT RiboGreen RNA Assay Kit (#R11490, Invitrogen) or a NanoDrop spectrophotometer, and RNA integrity was assessed on an Agilent Bioanalyzer 2100 system, for tissue, RiboTag and CD11b⁺ samples, a RIN > 7 was considered suitable for RNA-seq. Sample purity was verified by qRT-PCR.

### Quantitative real-time PCR

Total RNA was reverse transcribed and amplified in a one-step reaction using the Power SYBR Green RNA-to-CT 1-Step Kit (#4389986, Applied Biosystems) or the TaqMan RNA-to-CT 1-Step Kit (#4392938, Applied Biosystems), with equal RNA amounts per reaction, on an AriaMx Real-Time PCR System (Agilent). Relative expression levels were determined by the standard curve method (Sanz *et al*., 2019). Amplification efficiency (80-120%) and linearity (r² ≥ 0.99) were verified for all assays. TaqMan probes and SYBR Green primer pairs are listed in Table S2.

### RNA-seq and bioinformatic analysis

For profiling of *Gad2*- and *Vglut2-Ndufs4*-deficient neurons, mRNA libraries were prepared in-house from total RNA using the NEBNext Poly(A) mRNA Magnetic Isolation Module (#E7490S, New England Biolabs) and the NEBNext Ultra II Directional RNA Library Prep Kit for Illumina (#E7765S, New England Biolabs), with indices from the NEBNext Multiplex Oligos for Illumina Set 1 (#E7335, New England Biolabs). Library size and quality were verified on a Bioanalyzer 2100 (DNA 1000 kit, #5067-1504, Agilent). Sequencing was performed at the EMBL Genomics Core Facility (Heidelberg, Germany) on an Illumina NextSeq500 platform with paired-end (PE) sequencing at a read length of 2×75 base pairs and a sequencing depth of ∼30 million reads per sample. Bioinformatic analysis was conducted in-house: raw fastq files were quality-checked with FastQC and trimmed from base 7 to 77 with FastX Trimmer, reads were aligned to the mouse reference genome (Mus_musculus.GRCm38, Ensembl annotation) using STAR, and gene-level counts were obtained with HTSeq-count (minimum alignment quality, 4). Differential expression analysis was conducted using the DESeq2 (version 1.22.2) package (Love et al., 2014), considering transcripts with an adjusted p-value (padj) < 0.05 as differentially expressed. Gene ontology (GO) analyses were performed using the WebGestalt online tool.

For CD11b⁺ cells and Ifnar1DKO experiments, library preparation, sequencing and bioinformatic analysis were performed at Biomarker Technologies (BMK) GmbH (Münster, Germany). Low-input mRNA libraries were prepared using the NEBNext Poly(A) mRNA Magnetic Isolation Module (#E7490) and the NEBNext Ultra II Directional RNA Library Prep Kit for Illumina (#E7760L). Sequencing was performed on an Illumina NovaSeq6000 platform (paired-end, 2 × 150 bp, 20 million reads per sample). Differential expression was analysed with DESeq2 (version 1.30.1) (Love *et al*., 2014), considering transcripts with FC > 1.5 and an adjusted p-value below 0.05, and GO analyses were performed with WebGestalt.

### Brain organoids RNA-seq

Bulk RNA-seq of striatal brain organoids was conducted for WT and NDUFS4_1 samples. For each genotype, five organoids were pooled into a single pellet and snap-frozen, with each pellet obtained from an independent differentiation batch (n=2 independent experiments per genotype). Total RNA was isolated using the Qiagen isolation kit and quality checked by Nanodrop analysis (Nanodrop Technologies). Briefly, total RNA was mixed with 1 μg of a DNA oligonucleotide pool comprising 50-nt long oligonucleotide mix covering the reverse complement of the entire length of each rRNA (28S rRNA, 18S rRNA, 16S rRNA, 5.8S rRNA, 5S rRNA, 12S rRNA), incubated with 1U of RNase H (Hybridase Thermostable RNase H, Epicentre), purified using RNA Cleanup XP beads (Agencourt), DNase treated using TURBO DNase rigorous treatment protocol (Thermo Fisher Scientific, USA) and purified again with RNA Cleanup XP beads. rRNA-depleted RNA samples were further fragmented and processed into strand-specific cDNA libraries using TruSeq Stranded Total LT Sample Prep Kit (Illumina, USA) and sequenced on NextSeq 500, High Output Kit, 1 × 150 cycles. Raw RNA-seq data were processed to generate FASTQ files using Illumina bcl2fastq. Briefly, base calling was performed using Real-Time Analysis (RTA) software, and adapter sequences were trimmed. The resulting BCL files were converted to FASTQ format, followed by quality control assessment using FastQC before downstream analysis. The preprocessing of the FASTQ raw sequencing files was performed with nfcore/rnaseq (version 3.12.0) pipeline. Briefly, reads were trimmed with Trim Galore! (version 0.6.10) and mapped to the human genome (GRCh38 assembly) using STAR (version 2.7.10a) aligner. Transcript counts were determined using Salmon (version 1.10.1). DEGs were identified with DESeq2 (version 1.40.2) (Love *et al*., 2014), considering transcripts with a Benjamini– Hochberg-adjusted p-value below 0.05. GO analyses were performed with WebGestalt.

### PKR IP and dsRNAseq

Pools of frozen OBs from Ndufs4KO and control mice (Late stage; n=3 pools of 3 animals each) were dissected on ice, minced in ice-cold PBS and collected by centrifugation for 30 s at 4°C. Protein-RNA complexes were crosslinked in 750 μL of 0.1% PFA in PBS for 10 min at RT with rotation, and the reaction was quenched with 250 μL of 1 M glycine for 10 min. After two washes in PBS, pellets were resuspended in 400 μL of fCLIP lysis buffer supplemented with 1× protease inhibitor cocktail (#P8340, Sigma-Aldrich) and RNasin Plus RNase Inhibitor (1:100, #N2615, Promega). Samples were homogenized with a Polytron for 30 s (#10283722, Kinematica), incubated on ice for 10 min and sonicated for 2 min (30 s on/off cycles, 30% amplitude). NaCl was added to a final concentration of 150 mM and lysates were cleared at 13,000xg for 10 min at 4°C, keeping a 40 μL aliquot as input. PKR was immunoprecipitated overnight at 4°C with rotation using an anti-PKR antibody (#ab184257, Abcam). Complexes were captured on Protein A/G Magnetic Beads (#88803, Pierce) for 3 h at 4°C, washed and eluted in 300 μL of fCLIP elution buffer for 3 h at 25°C. Finally, crosslinks were reversed and proteins digested by adding proteinase K (20 mg/mL) and incubating overnight at 65 °C.

RNA was extracted with acid phenol-chloroform (1 h at 37°C, then 12,000xg for 10 min at RT) and precipitated overnight at −20°C with 1/10 volume of 3 M sodium acetate, glycogen (20 μg/μL) and one volume of isopropanol. Pellets were recovered by centrifugation at maximum speed for 1 h at 4°C, washed twice with 500 μL of 75% ethanol (12,000 xg, 5 min, 4°C) and air-dried at RT. Residual DNA was removed by incubating each sample for 15 min at RT with DNase I in RDD buffer (RNase-Free DNase Set, #79254, Qiagen) supplemented with RNase inhibitor. RNA was then re-extracted with acid phenol-chloroform, precipitated overnight at −80°C and resuspended in 10 μL of nuclease-free water. RNA-seq libraries were prepared from PKR-immunoprecipitated RNA at GenXPro (Frankfurt, Germany) using the NEBNext Ultra II Directional RNA Library Prep Kit (#E7760, New England Biolabs) according to the manufacturer’s instructions. Briefly, total RNA was fragmented, followed by cDNA synthesis using random hexamers. The resulting cDNA fragments underwent end repair and A-tailing followed by ligation of Illumina p5 and p7 adapters. After PCR amplification using a minimum number of cycles, libraries were purified with silica beads and the quality was controlled for size-distribution and quantity. Sequencing was performed on an Illumina NextSeq 500 instrument (single-end, 1 × 76 bp). Unprocessed sequencing reads were adapter-and quality-trimmed using Cutadapt (3.4), and read quality was assessed with FastQC (v0.11.9). Trimmed reads were mapped with STAR (v2.7.10a), allowing spliced alignments, to the Mus musculus reference genome (GRCm39, Ensembl release 105), which includes the mitochondrial genome. Quantification of transcript abundance was performed using RSEM (v1.3.3). Raw counts were geometric-mean-normalized using DESeq2 (v1.38) and expressed as TPM. Differential expression analysis was performed with DESeq2 (v1.38). Quality metrics were aggregated across samples with MultiQC (v1.12). Strand-specific coverage was plotted separately for heavy- and light-strand-derived reads over the *mt-Nd1-mt-Nd6* interval (GRCm39), excluding rRNA genes from the plotted region.

### Statistical analyses

GraphPad Prism software (v8) was used for statistical analyses. Data are shown as the mean ± SEM. The values for n, p-value and the specific statistical test performed are indicated in each figure legend; n refers to the number of animals per group, except where samples were pooled, as specified in the corresponding legend. *P < 0.05, **P < 0.01, ***P < 0.001 and ****P < 0.0001.

## Data availability

RNA-seq datasets generated in this study have been deposited in the NCBI Sequence Read Archive under BioProject accessions PRJNA1513771 (RiboTag translatome of OB GABAergic neurons), PRJNA1534213 (RiboTag translatome of vestibular glutamatergic neurons), PRJNA1502843 (CD11b⁺ cells from *Ndufs4*-deficient mouse models) and PRJNA1553472 (PKR-bound RNA sequencing), and brain organoid RNA-seq data. Mass spectrometry data and search result files have been deposited in the ProteomeXchange Consortium via the jPOST partner repository (https://repository.jpostdb.org)(Okuda et al., 2017) under the identifiers PXD086295 (ProteomeXchange) and JPST005032 (jPOST).

## Author contributions

MGT designed research studies, conducted experiments, acquired data, analyzed data, and wrote the manuscript. PB and MBR designed research studies, conducted experiments, acquired data, and analyzed data. MLS, PPD, EP, CC, LC, AG, MG, LSB, and BA conducted experiments, acquired data, and analyzed data. JFI, ESa, MA, and LF acquired and analyzed proteomic and sequencing data. SL, PL, AdS and AP generated and characterized human iPSC-derived organoids and analyzed data. GM generated STING1DKO mice and analyzed data. ES and AQ designed research studies, analyzed data, provided reagents, and wrote the manuscript. All authors reviewed and approved the final version of the manuscript.

## Acknowledgments

The authors thank all the lab members for their feedback; Dr. Adolfo García-Sastre (Icahn School of Medicine at Mount Sinai, USA) for providing *Eif2ak2*-deficient mice; Dr. Ole Kiehn (Karolinska Institutet, Sweden) for *Slc17a6*-Cre mice; Prof. Richard Palmiter (University of Washington, USA) for the pAAV-CRE·GFP and pAAV-ΔCRE·GFP plasmids; Prof. Kobi Rosenblum (University of Haifa, Israel) for the PKR shRNA sequence.

This work was supported by three MINECO Ramón y Cajal fellowships (RYC2020-029596-I, EP; RYC2019-028501-I, ES; and RYC-2012-11873, AQ), a MICINN Ramón y Cajal fellowship (RYC2022-037332-I, LC), Marie Skłodowska-Curie fellowships (H2020-MSCA-IF-2020-101030783, MLS; H2020-MSCA-COFUND-2014-665919, AG), and predoctoral fellowships from MICIU (PRE2018-083179 to LSB; FPU19/03990 to MGT) and from AGAUR, Generalitat de Catalunya (2023FI-100663 to MBR). EP received funds from MICINN (PID2021-125079OA-I00, PID2024-161421OB-I00 and CNS2024-154890), and from the SynLeigh project (ERDERA25/17), funded by the Instituto de Salud Carlos III (#AC25/00124), a funding partner of the European Rare Diseases Research Alliance (ERDERA), and co-funded by the European Union under the Horizon Europe programme (Grant Agreement No 101156595 - ERDERA). ES received funds from MICINN (PID2019-107633RB-I00 and PID2022-142544OB-I00) and MICIU (CNS2025-166925 and PID2025-175346NB-I00). AQ is a recipient of an ICREA Academia award and received funds from the European Research Council (Starting Grant NEUROMITO, ERC-2014-StG-638106), PID2020-114977RB-I00, PID2023-151947OB-I00, RED2022-134786-T and RED2024-154130-T, funded by MICIU/AEI/10.13039/501100011033, by ERDF “A way of making Europe” and by NextGenerationEU/PRTR, AGAUR (2017SGR-323, 2021SGR-720), Fundació TV3-La Marató (202030), 415/C/2020, and “la Caixa” Foundation (ID 100010434, under the agreements LCF/PR/HR20/52400018 and LCF/PR/HM25/49010001). AP receives funds from the Deutsche Forschungsgemeinschaft (DFG) (PR1527/13-1, PR1527/14-1, PR1527/15-1, PR1527/18-1 to AP), ERDERA and the Federal Ministry of Research, Technology and Space (BMFTR) (SynLeigh #01GM2602A to AP), the European Commission’s Horizon Europe Programme (SIMPATHIC #101080249 to AP) and AFM-Téléthon (28545, 29044, 30119, to AP).

## Supplementary material

**Figure S1.**
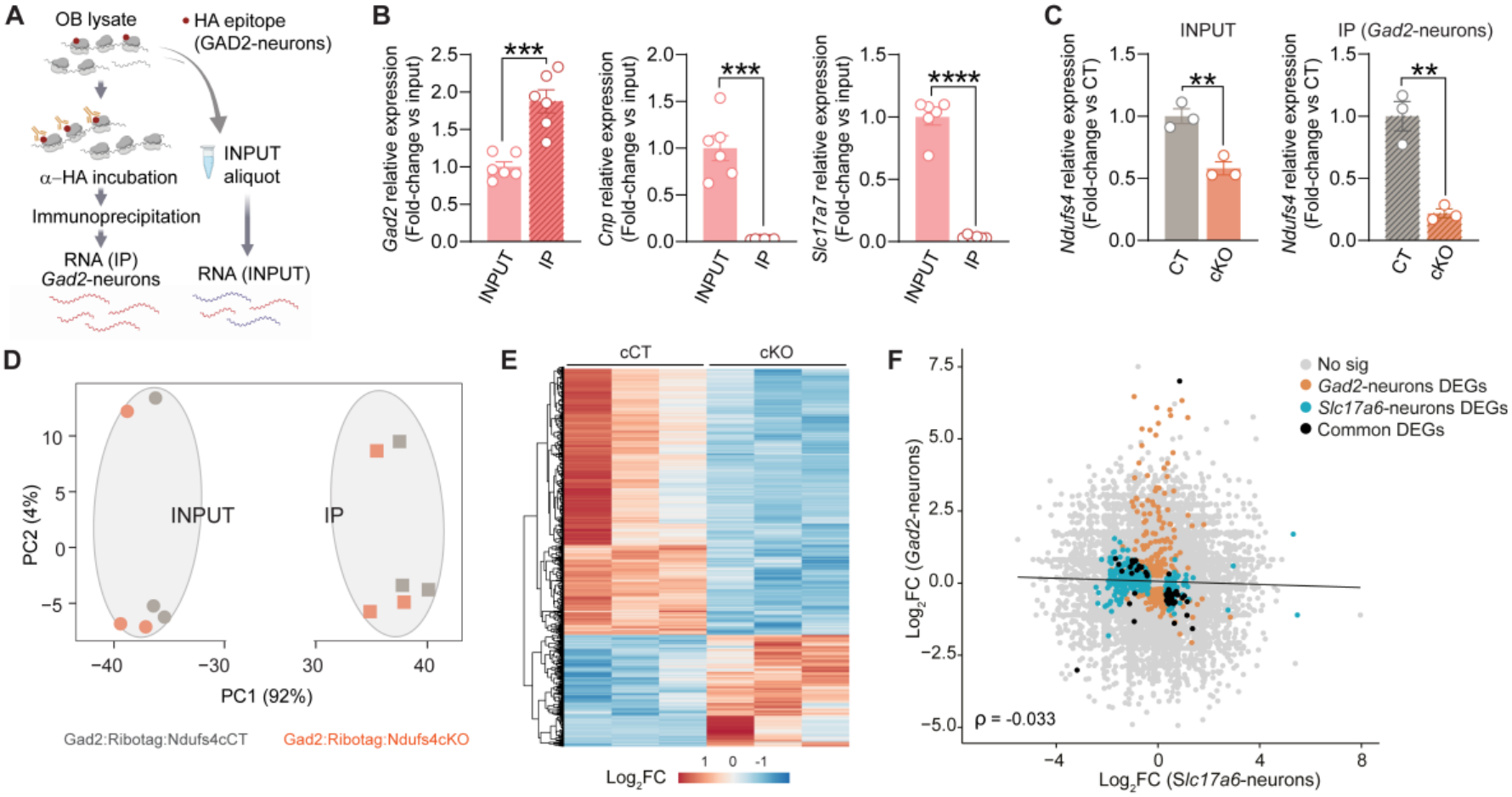
Cell-type specific translatome profiling of *Ndufs4*-deficient neurons. (A) Schematic of the RiboTag approach: HA-tagged ribosomes from *Gad2⁺* (GAD2) neurons are immunoprecipitated (α-HA) to isolate neuron-specific mRNA (IP), keeping a whole-tissue aliquot as INPUT. (B) qRT-PCR validation showing enrichment of the GABAergic marker *Gad2* in the IP fraction and depletion of the non-target markers *Cnp* (oligodendrocytes) and *Slc17a7* (glutamatergic neurons) (n=6). Mean ± SEM; unpaired Student’s t-test. ***p < 0.001, ****p < 0.0001. (C) qRT-PCR of *Ndufs4* in INPUT and IP fractions from Gad2:RiboTag:Ndufs4cCT (CT) and Gad2:RiboTag:Ndufs4cKO (cKO) mice (n=3). Mean ± SEM; unpaired Student’s t-test. **p < 0.01. (D) PCA of INPUT and IP RNA-seq samples from Gad2:RiboTag:Ndufs4cCT (CT) and Gad2:RiboTag:Ndufs4cKO (cKO) mice; PC1 (92% of variance) separates IP from INPUT fractions. (E) Heatmap of differentially expressed genes in IP samples (cKO vs CT). (F) Spearman correlation between transcriptional responses to *Ndufs4* deletion in glutamatergic and GABAergic neurons. Each dot represents a transcript; axes show Log_2_FC (cKO vs CT) in the IP fraction of vestibular *Slc17a6-*expressing neurons (Vglut2:RiboTag:Ndufs4cKO; n=3) and OB *Gad2-*expressing neurons (Gad2:RiboTag: Ndufs4cKO; n=3). Transcripts differentially expressed (DEGs) only in *Gad2*⁺ neurons (orange), only in *Slc17a6*⁺ neurons (cyan), in both (black) or in neither (grey) are indicated (DESeq2 adjusted p<0.05). Spearman’s ρ=-0.033. Line shown for visualization only.

**Figure S2.**
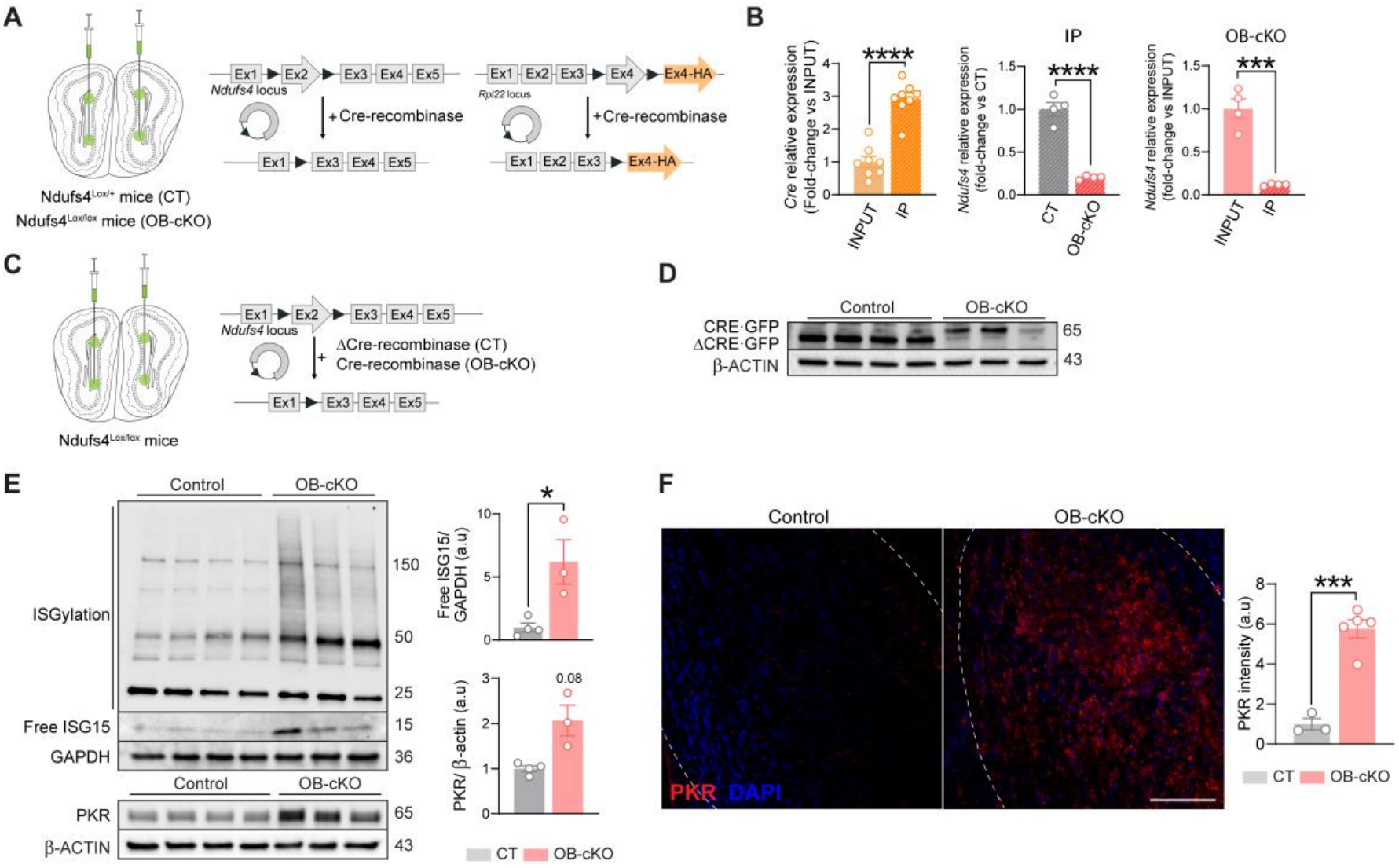
Targeted *Ndufs4* deletion is sufficient to trigger antiviral signaling. (A) Schematic of AAV1-mediated *Ndufs4* deletion and ribosome HA-tagging in the OB of Ndufs4^lox/+^Rpl22^HA/HA^ (control) and Ndufs4^lox/lox^Rpl22^HA/HA^ (OB-cKO) mice (B) qRT-PCR validation: *Cre* enrichment in the IP vs INPUT fraction (n=8), *Ndufs4* levels in the IP fraction of CT vs OB-cKO mice (n=4), and *Ndufs4* levels in IP vs INPUT within OB-cKO mice (n=4). Mean ± SEM; unpaired Student’s t-test. ***p < 0.001, ****p < 0.0001. (C) Schematic of the ΔCRE-injected (control) vs CRE-injected (OB-cKO) Ndufs4^lox/lox^ design. (D) Immunoblot of CRE·GFP and ΔCRE·GFP confirming transduction (n=3-4). (E) Immunoblot and quantification of the antiviral proteins ISG15 and PKR in CRE-(OB-cKO) vs ΔCRE-injected (CT) mice relative to β-ACTIN or GAPDH (n=3-4). Mean ± SEM; unpaired t-test for Free ISG15 and unpaired t-test with Welch’s correction for PKR. *p<0.05. (F) Representative immunofluorescence and quantification of PKR intensity in the OB of CT and OB-cKO mice (n=3-5). Scale bar = 150 µm; Mean ± SEM; unpaired t-test. ***p < 0.001.

**Figure S3.**
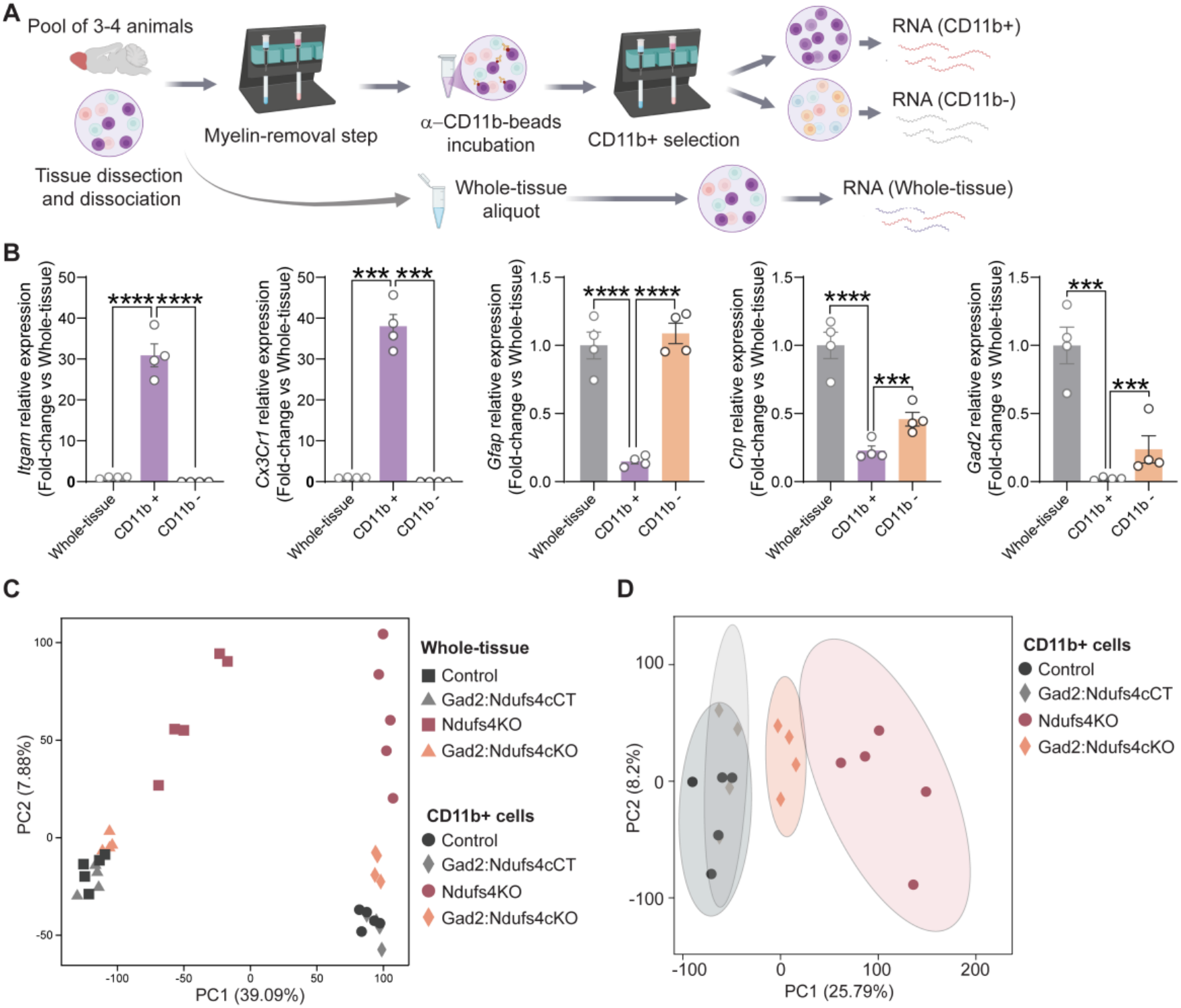
Isolation and transcriptomic profiling of OB CD11b+ cells. (A) Schematic of magnetic-bead isolation of CD11b+ cells, yielding RNA from CD11b⁺, CD11b⁻ and whole-tissue fractions (pool of 3-4 animals per sample). (B) qRT-PCR of the microglial markers *Itgam* and *Cx3cr1* and of markers of other cell types (*Gfap, Cnp, Gad2*) in CD11b⁺ and CD11b⁻ fractions relative to whole tissue (n=4 pools). Mean ± SEM; one-way ANOVA followed by Tukey’s post hoc. ***p < 0.001, ****p < 0.0001. (C) PCA of whole-tissue and CD11b+ RNA-seq samples (Control, Gad2:Ndufs4cCT, Ndufs4KO, Gad2:Ndufs4cKO). (D) PCA restricted to CD11b⁺ samples (n=4-5 pools of 3-4 animals each).

**Figure S4.**
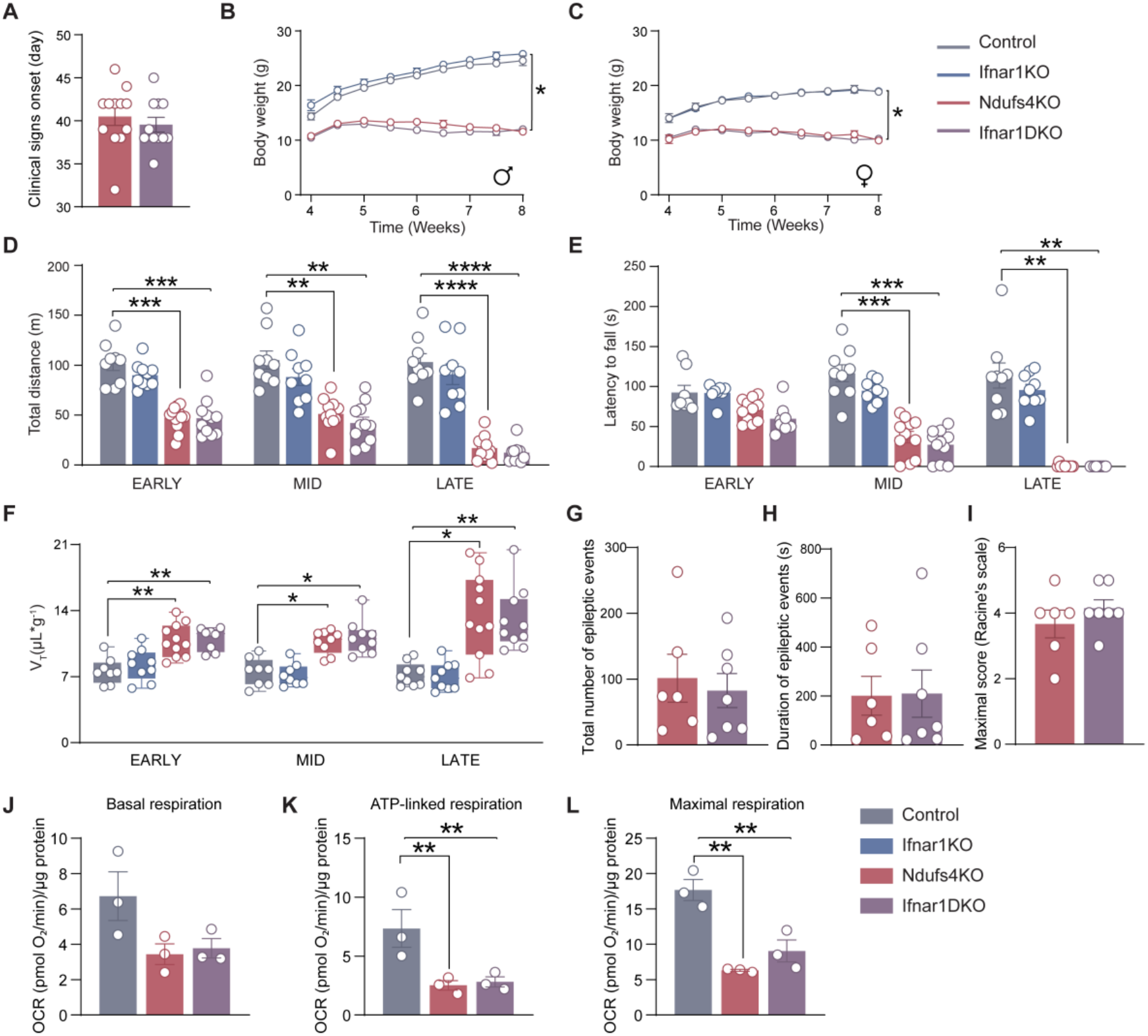
Effects of blocking IFN-I signaling on the Ndufs4KO phenotype. (A) Onset of clinical signs in Ndufs4KO (n=12) and Ifnar1DKO (n=11) mice. Mean ± SEM; unpaired Student’s t-test; ns. (B,C) Body weight curves of (B) male (n=7-13) and (C) female (n=6-12) mice of all four genotypes. Mean ± SEM; mixed-effects model. *p<0.05. (D) Total distance travelled in the open field at early, mid and late disease stages (n=9-12). Mean ± SEM; three-way ANOVA with Geisser-Greenhouse correction followed by Tukey’s post hoc. **p < 0.01, ***p < 0.001, ****p < 0.0001. (E) Latency to fall in the accelerating rotarod (n=8-12). Same analysis. **p < 0.01, ***p < 0.001. (F) Tidal volume (V_T_) by whole-body plethysmography (n=8-11). Box-and-whisker plots; same analysis. *p < 0.05, **p < 0.01. (G-I) Heat-induced seizure susceptibility at mid disease stage (PND42) in Ndufs4KO and Ifnar1DKO mice: (G) total number of epileptic events; (H) total duration (s); (I) maximal severity (modified Racine scale). Mean ± SEM (n=6-7); unpaired Student’s t-test; ns. (J-L) Oxygen consumption rate of mitochondria isolated from the OB of control, Ndufs4KO and Ifnar1DKO mice (n=3). Mean ± SEM; One-way ANOVA followed by Tukey’s post hoc. *p < 0.05, **p < 0.01.

**Figure S5.**
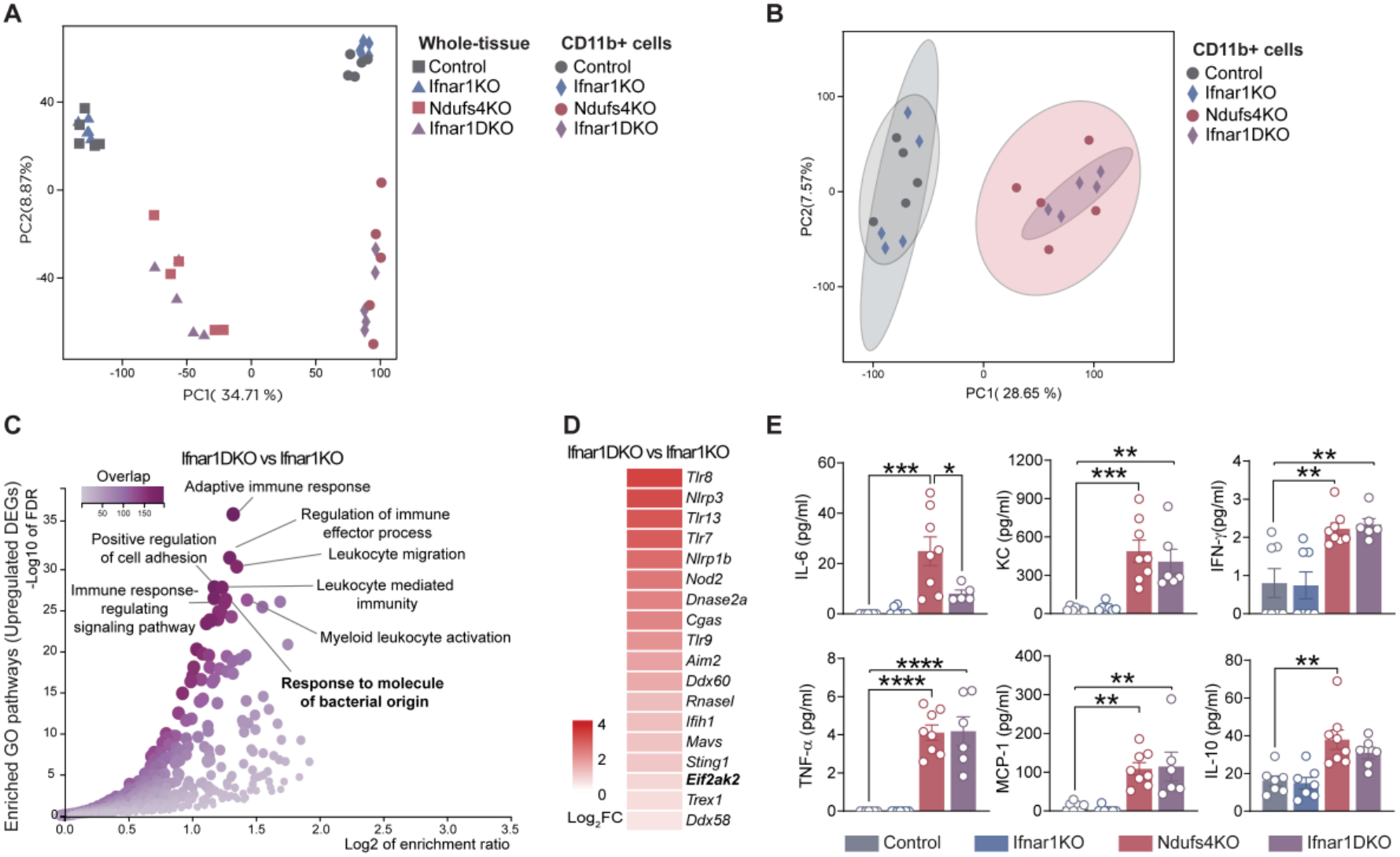
Molecular effects of blocking IFN-I signaling in Ndufs4KO mice. (A) PCA of whole-tissue and CD11b⁺ RNA-seq samples from control, Ifnar1KO, Ndufs4KO and Ifnar1DKO mice (n=5 biological replicates; each pooled from 3-4 animals). (B) PCA restricted to CD11b⁺ samples. (C) GO enrichment of upregulated pathways in whole-tissue OB samples from Ifnar1DKO vs Ifnar1KO mice. (D) Heatmap (Log_2_FC) of genes encoding innate immune sensors of nucleic acids in whole-tissue OB samples, Ifnar1DKO vs Ifnar1KO. (E) Cytokine and chemokine levels in OB samples measured by multiplex assay (n=5-8): IL-6, KC (CXCL1), IFN-γ, TNF-α, MCP-1 (CCL2) and IL-10. Mean ± SEM; two-way ANOVA followed by Tukey’s post hoc. *p < 0.05, **p < 0.01, ***p < 0.001, ****p < 0.0001.

**Figure S6.**
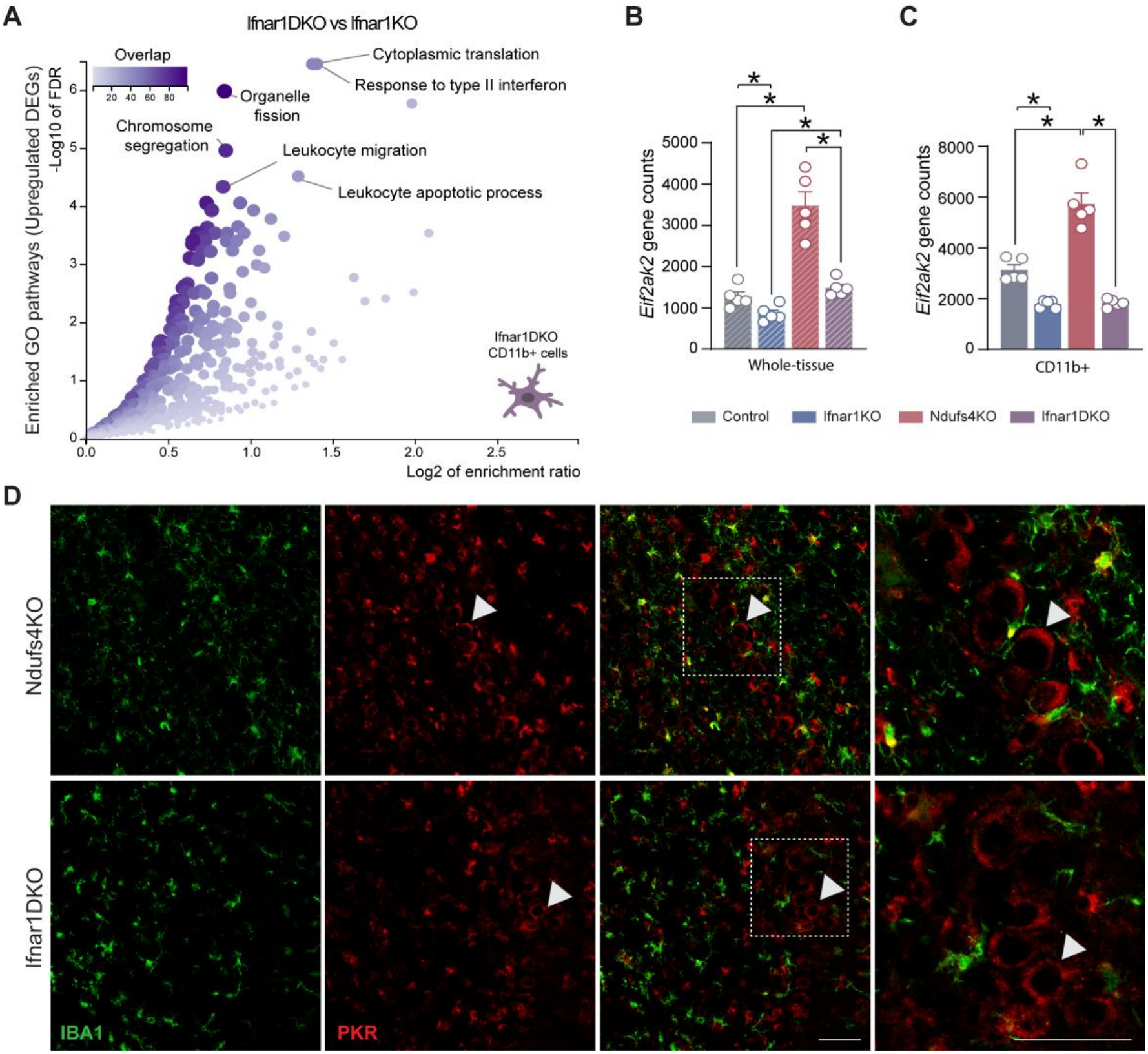
PKR is predominantly induced in cell types other than microglia in *Ifnar1*-deficient Ndufs4KO mice. (A) GO enrichment of upregulated pathways in CD11b⁺ cells from Ifnar1DKO vs Ifnar1KO mice showing the absence of antiviral/IFN-I GO terms. (B-C) *Eif2ak2* normalized gene counts in whole-tissue (B) and CD11b⁺ (C) RNA-seq samples from all four genotypes (n=5 biological replicates; each pooled from 3-4 animals). Mean ± SEM; Statistics refer to the DESeq2 pairwise comparisons, which correct for multiple comparisons. *p < 0.05. (D) IBA1 (green) and PKR (red) immunostaining in the OB of Ndufs4KO and Ifnar1DKO mice; dashed boxes indicate the magnified regions and arrowheads indicate representative PKR⁺/IBA1⁻ cells. Scale bars, 50 µm.

**Figure S7.**
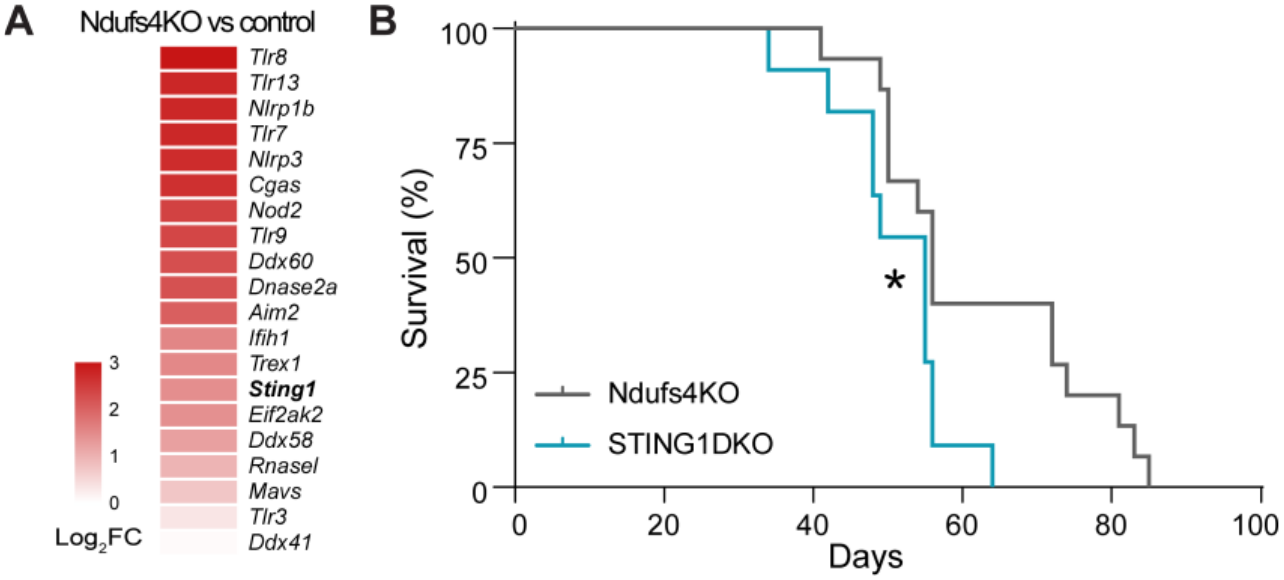
STING1 deletion does not improve survival in Ndufs4KO mice. (A) Heatmap (Log_2_FC) of genes encoding innate immune sensors of nucleic acids in whole-tissue OB samples, Ndufs4KO vs CT. (B) Survival curves of Ndufs4KO (n=15) and STING1DKO (n=11) mice. Log-rank test; *p< 0.05

**Figure S8.**
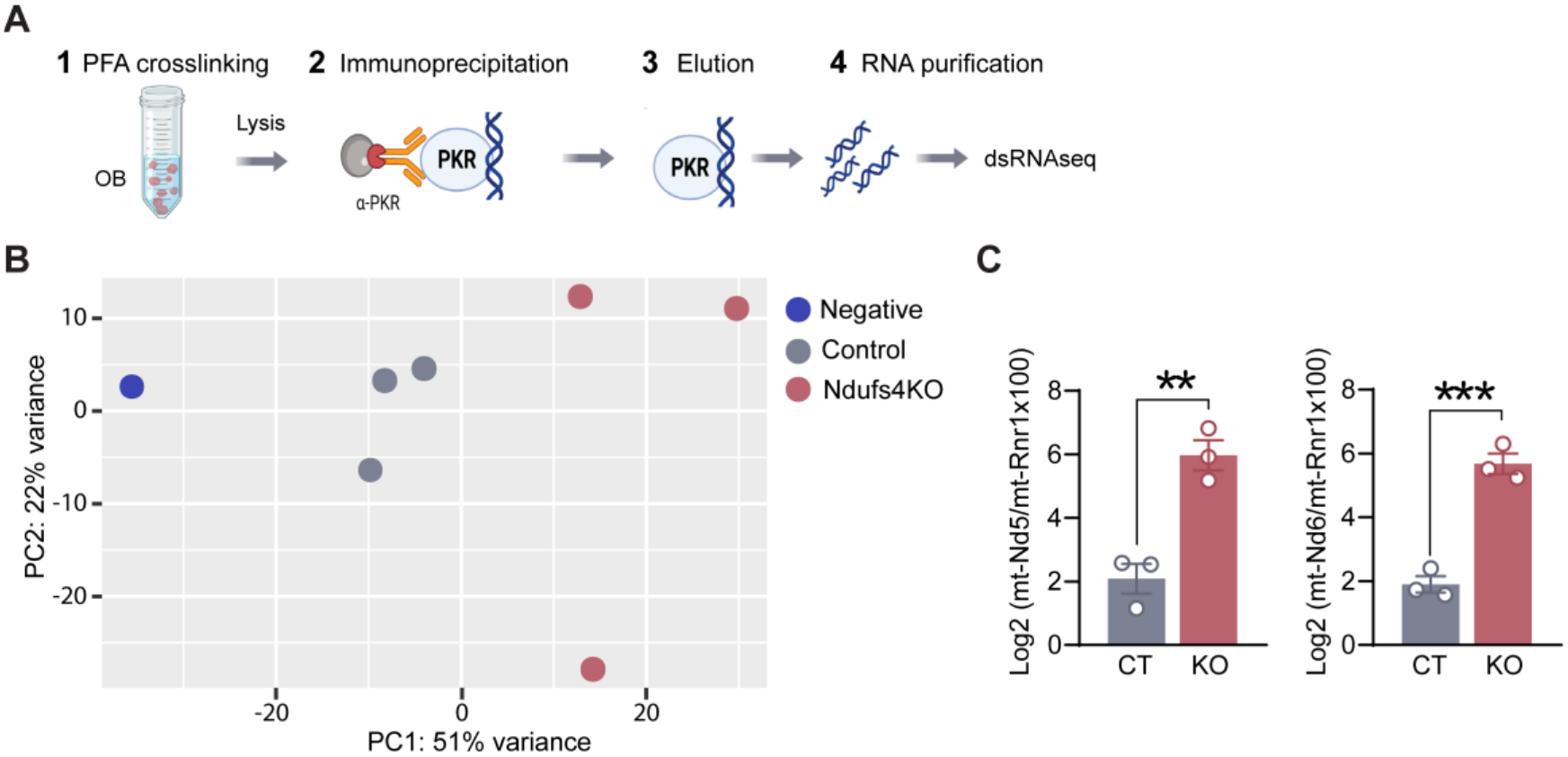
PKR-dsRNA sequencing. (A) Schematic of the PKR fCLIP and dsRNA sequencing workflow. (B) Principal component analysis of the PKR-bound transcriptomes of control and Ndufs4KO samples (n=3 pools of 3 animals each), together with a mock immunoprecipitation performed with an anti-GFP antibody (Negative, n=1). (C) Relative abundance of the duplex-forming transcripts *mt-Nd5* and *mt-Nd6* in PKR IPs from control (CT) and Ndufs4KO (KO) mice (n=3 pools per group). FPKM values are expressed relative to *mt-Rnr1*, a mitochondrial transcript outside the duplex-forming region, to report enrichment relative to overall mitochondrial RNA content, and shown as Log_2_ (ratio × 100). Mean ± SEM; unpaired Student’s t-test on Log_2_-transformed values. **p < 0.01, ***p < 0.001.

**Figure S9.**
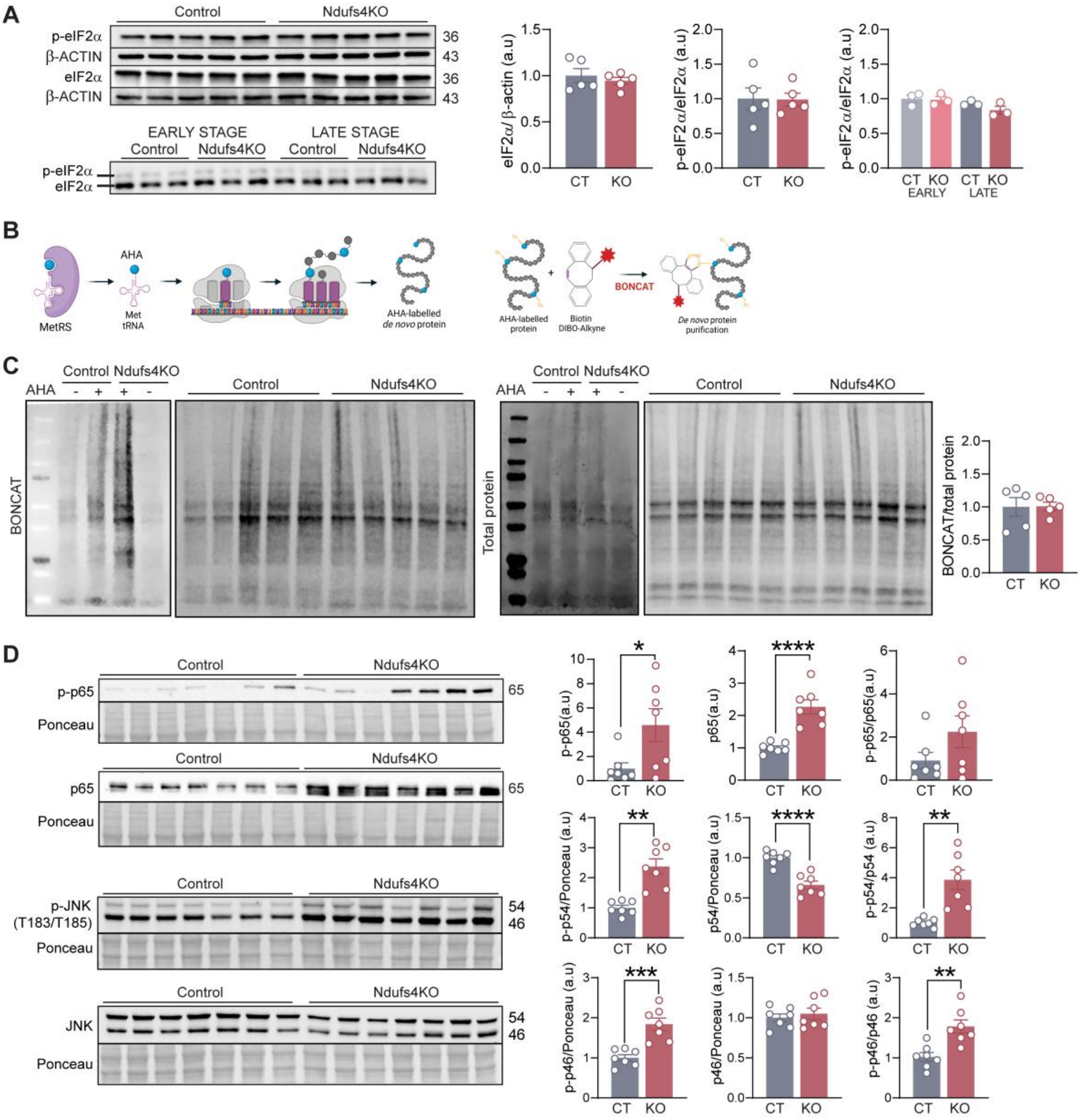
PKR activation does not engage global translational repression but is associated with proinflammatory signaling. (A) Representative immunoblots and quantification of total eIF2α and phospho-eIF2α (Ser51) by conventional SDS-PAGE (n=5, late stage) and of the p-eIF2α/eIF2α ratio by Phos-tag SDS-PAGE at early and late disease stages (n=3). Mean ± SEM; unpaired Student’s t-test or two-way ANOVA; ns. (B) Diagram depicting visualization of newly synthesized proteins via AHA labelling. Using the endogenous translational machinery, de novo synthesized proteins are labelled with AHA at their amino-terminal and internal methionine residues and covalently bonded to a tag through interaction of the azide group of AHA with the alkyne group of the tag, enabling purification by bio-orthogonal non-canonical amino acid tagging (BONCAT). Adapted from (Evans *et al*., 2020). (C) Representative BONCAT immunoblot and total-protein loading control, including AHA-negative controls, with quantification of AHA-labelled protein normalized to total protein (n=5). Mean ± SEM; unpaired Student’s t-test; ns. (D) Representative immunoblots and quantification of phospho-p65 (Ser536), total p65 and the p-p65/p65 ratio, and of phospho-JNK and total JNK, with the p-p54/p54 and p-p46/p46 ratios, in the OB of control and Ndufs4KO mice, normalized to Ponceau S (n=7). Mean ± SEM; unpaired Student’s t-test with Welch’s correction for p-p65, phospho-p54 and the p-p54/p54 ratio and unpaired Student’s t-test for the remaining comparisons. *p < 0.05, **p < 0.01, ****p < 0.0001.

**Figure S10.**
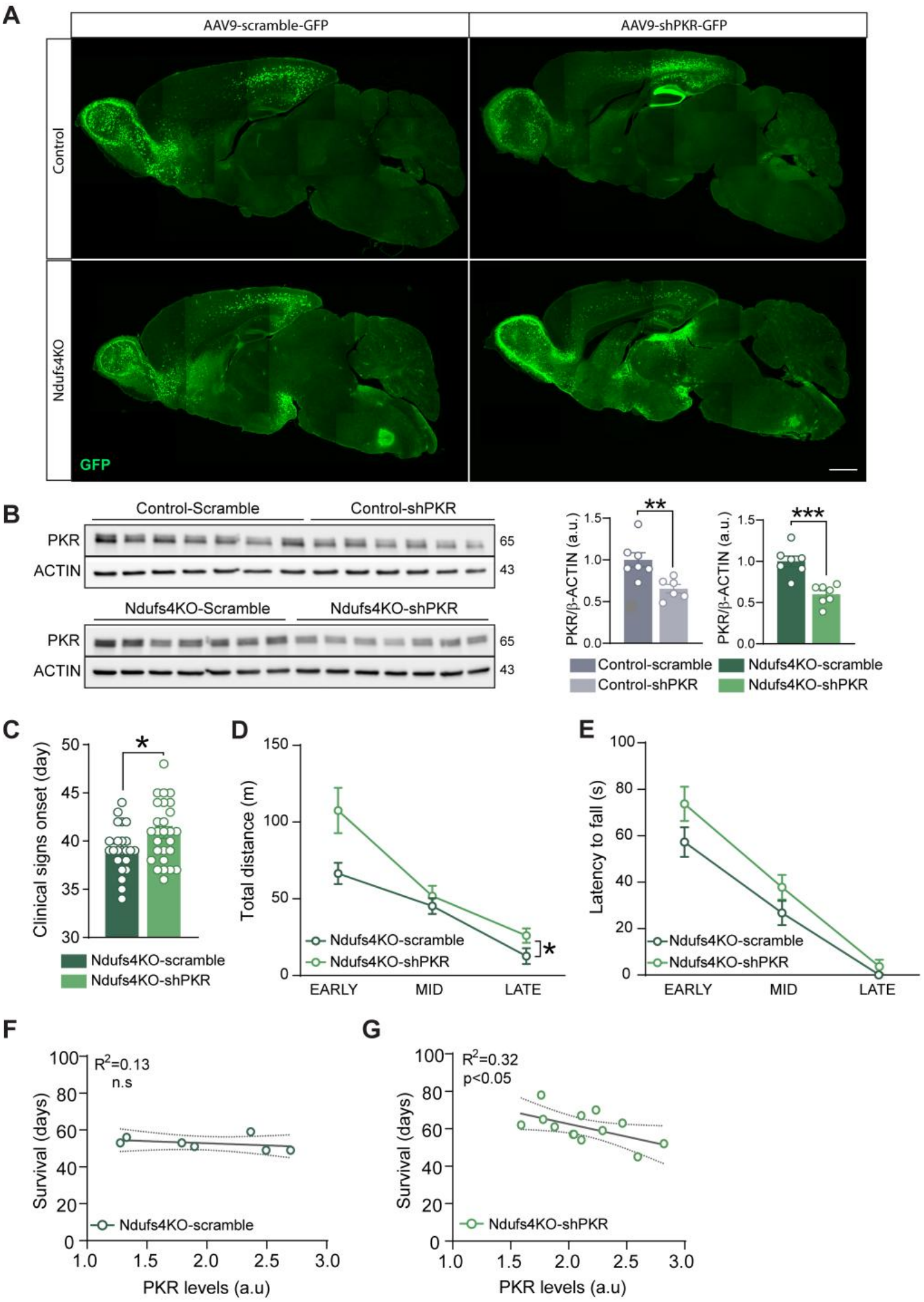
PKR silencing by AAV double-administration in Ndufs4KO mice. (A) Representative image of GFP signal in sagittal brain sections from control and Ndufs4KO mice injected with AAV9-scramble-GFP or AAV9-shPKR-GFP. Scale bar, 1 mm. (B) Representative immunoblots of PKR in OB samples from scramble-and shPKR-injected control (upper) and Ndufs4KO (lower) mice, with quantification normalized to β-ACTIN (n=6-7). Mean ± SEM; unpaired Student’s t-test. **p < 0.01, ***p < 0.001. (C) Onset of clinical signs in scramble-(n=23) and shPKR-injected (n=27) Ndufs4KO mice. Mean ± SEM; unpaired Student’s t-test. *p < 0.05. (D) Spontaneous locomotion (total distance) in the open field in scramble-and shPKR-injected Ndufs4KO mice. Mean ± SEM; two-way ANOVA (mixed model) with Geisser-Greenhouse correction. *p < 0.05. (E) Latency to fall in the accelerating rotarod in scramble- and shPKR-injected Ndufs4KO mice (n=8-16). Same analysis; p = 0.1. (F,G) Correlation between survival and PKR protein levels (measured by western blot) in (F) scramble-injected (n=7) and (G) shPKR-injected Ndufs4KO mice (n=12). Linear regression; n.s for scramble; p = 0.04 for shPKR.

**Figure S11.**
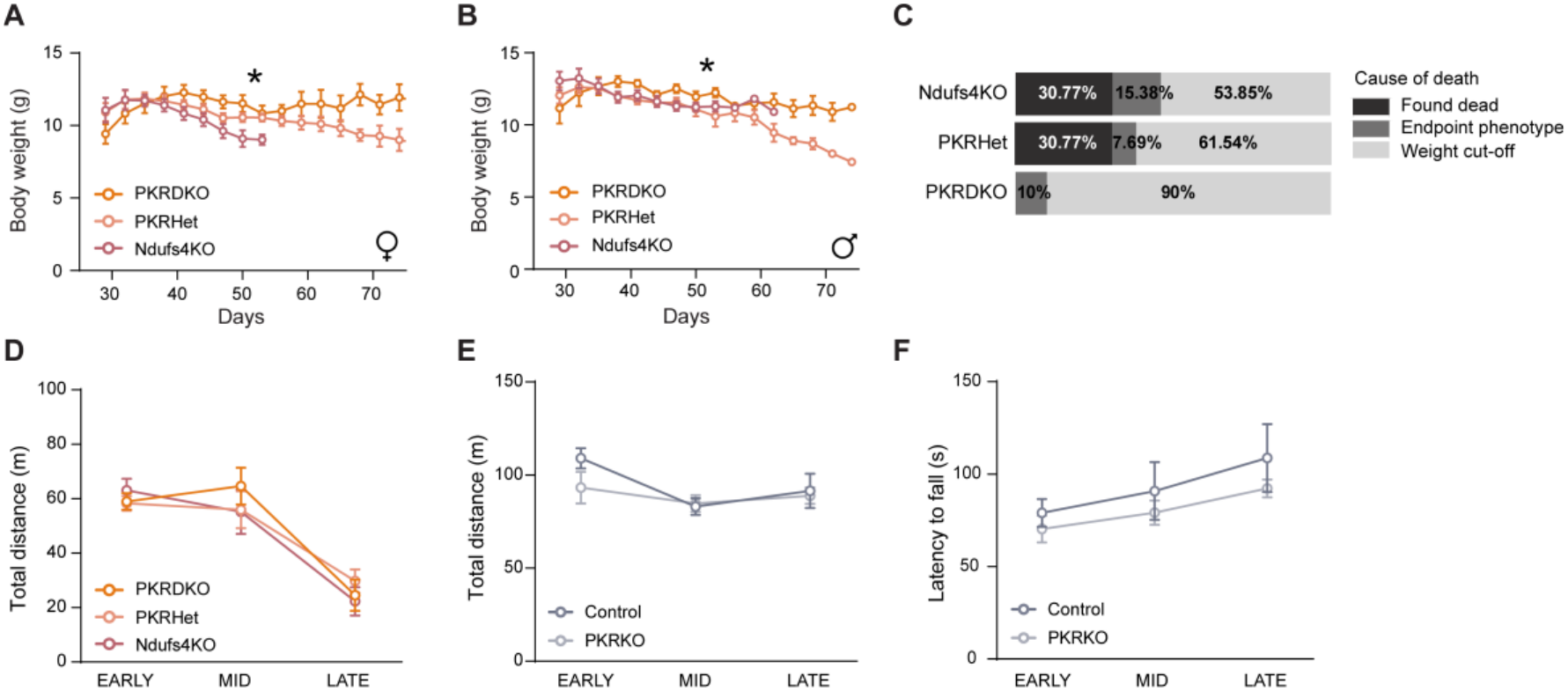
Effects of genetic PKR deletion on the Ndufs4KO phenotype. (A,B) Body weight curves of (A) female and (B) male Ndufs4KO, PKRHet and PKRDKO mice. Mean ± SEM; mixed-effects model. *p < 0.05 (n=8-9). (C) Cause of death across Ndufs4KO, PKRHet and PKRDKO mice (n=10-13). (D) Total distance in the open field in Ndufs4KO mice. Mean ± SEM; mixed-effects model (n=11-16). (E) Open field analysis in control and PKRKO mice. Mean ± SEM; mixed-effects model (n=8-11). (F) Latency to fall in the accelerating rotarod in control and PKRKO mice. Mean ± SEM; mixed-effects model (n=8-11).

**Figure S12.**
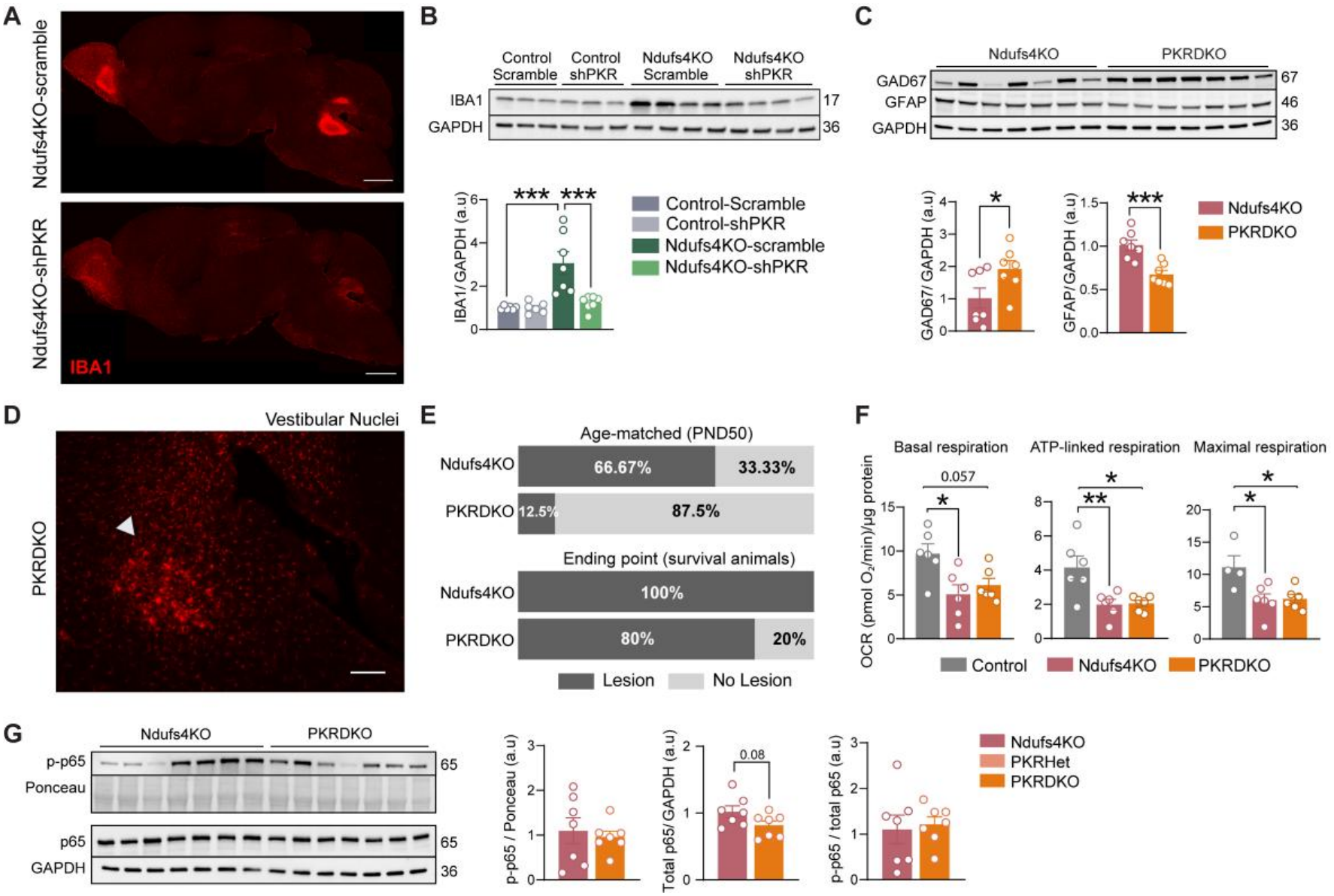
Impact of PKR silencing and deletion on the Ndufs4KO mice. (A) Representative immunofluorescence of IBA1 in sagittal brain sections of scramble- and shPKR-injected Ndufs4KO mice. Scale bar, 1 mm. (B) Representative immunoblot and quantification of IBA1 normalized to GAPDH in control and Ndufs4KO mice injected with scramble or shPKR (n=6-7). Mean ± SEM; two-way ANOVA followed by Tukey’s post hoc. ***p < 0.001. (C) Immunoblot and quantification of GAD67 and GFAP in the OB of Ndufs4KO and PKRDKO mice, normalized to GAPDH (n=7). Mean ± SEM; unpaired Student’s t-test. *p < 0.05, ***p < 0.001. (D) Representative image of VN lesion in PKRDKO survival animal. Scale bar = 100 µm. (E) Proportion of Ndufs4KO and PKRDKO mice presenting the characteristic vestibular nucleus lesion at age-matched (n=6-8) and at ending point (n=5). (F) Oxygen consumption rate of mitochondria isolated from the OB of control, Ndufs4KO and PKRDKO mice (n=5-6). Mean ± SEM; One-way ANOVA followed by Tukey’s post hoc. *p < 0.05, **p < 0.01. (G) Representative immunoblots and quantification of total p65, phospho-p65 (p-p65) and the p-p65/p65 ratio in the OB of Ndufs4KO and PKRDKO mice, normalized to GAPDH (n=7). Mean ± SEM; unpaired Student’s t-test.

---

**Table S1.**
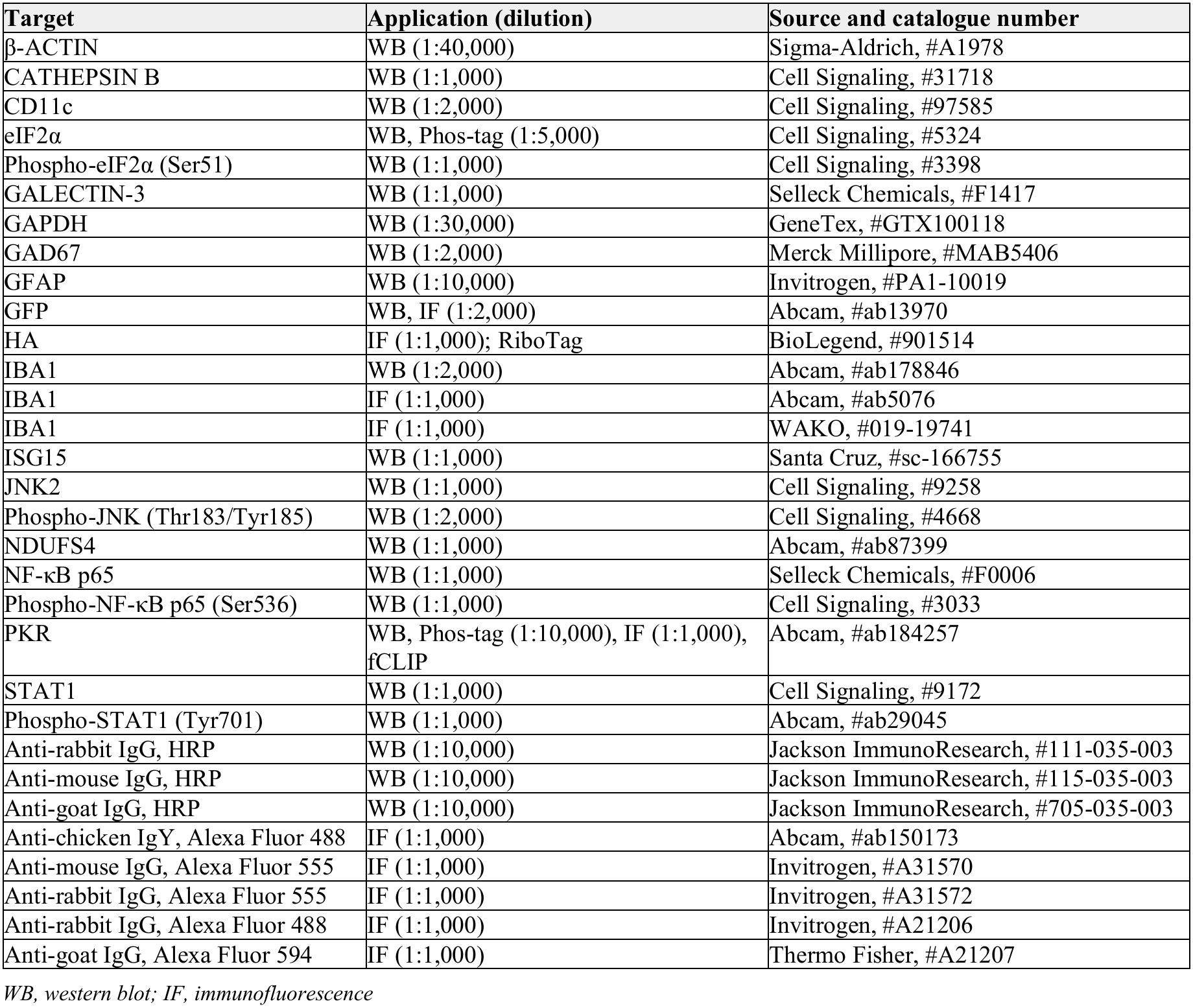
Antibodies used in this study.

| Target | Application (dilution) | Source and catalogue number |
|---|---|---|
| β-ACTIN | WB (1:40,000) | Sigma-Aldrich, #A1978 |
| CATHEPSIN B | WB (1:1,000) | Cell Signaling, #31718 |
| CD11c | WB (1:2,000) | Cell Signaling, #97585 |
| eIF2α | WB, Phos-tag (1:5,000) | Cell Signaling, #5324 |
| Phospho-eIF2α (Ser51) | WB (1:1,000) | Cell Signaling, #3398 |
| GALECTIN-3 | WB (1:1,000) | Selleck Chemicals, #F1417 |
| GAPDH | WB (1:30,000) | GeneTex, #GTX100118 |
| GAD67 | WB (1:2,000) | Merck Millipore, #MAB5406 |
| GFAP | WB (1:10,000) | Invitrogen, #PA1-10019 |
| GFP | WB, IF (1:2,000) | Abcam, #ab13970 |
| HA | IF (1:1,000); RiboTag | BioLegend, #901514 |
| IBA1 | WB (1:2,000) | Abcam, #ab178846 |
| IBA1 | IF (1:1,000) | Abcam, #ab5076 |
| IBA1 | IF (1:1,000) | WAKO, #019-19741 |
| ISG15 | WB (1:1,000) | Santa Cruz, #sc-166755 |
| JNK2 | WB (1:1,000) | Cell Signaling, #9258 |
| Phospho-JNK (Thr183/Tyr185) | WB (1:2,000) | Cell Signaling, #4668 |
| NDUFS4 | WB (1:1,000) | Abcam, #ab87399 |
| NF-κB p65 | WB (1:1,000) | Selleck Chemicals, #F0006 |
| Phospho-NF-κB p65 (Ser536) | WB (1:1,000) | Cell Signaling, #3033 |
| PKR | WB, Phos-tag (1:10,000), IF (1:1,000), fCLIP | Abcam, #ab184257 |
| STAT1 | WB (1:1,000) | Cell Signaling, #9172 |
| Phospho-STAT1 (Tyr701) | WB (1:1,000) | Abcam, #ab29045 |
| Anti-rabbit IgG, HRP | WB (1:10,000) | Jackson ImmunoResearch, #111-035-003 |
| Anti-mouse IgG, HRP | WB (1:10,000) | Jackson ImmunoResearch, #115-035-003 |
| Anti-goat IgG, HRP | WB (1:10,000) | Jackson ImmunoResearch, #705-035-003 |
| Anti-chicken IgY, Alexa Fluor 488 | IF (1:1,000) | Abcam, #ab150173 |
| Anti-mouse IgG, Alexa Fluor 555 | IF (1:1,000) | Invitrogen, #A31570 |
| Anti-rabbit IgG, Alexa Fluor 555 | IF (1:1,000) | Invitrogen, #A31572 |
| Anti-rabbit IgG, Alexa Fluor 488 | IF (1:1,000) | Invitrogen, #A21206 |
| Anti-goat IgG, Alexa Fluor 594 | IF (1:1,000) | Thermo Fisher, #A21207 |

*WB, western blot; IF, immunofluorescence*

**Table S2.**
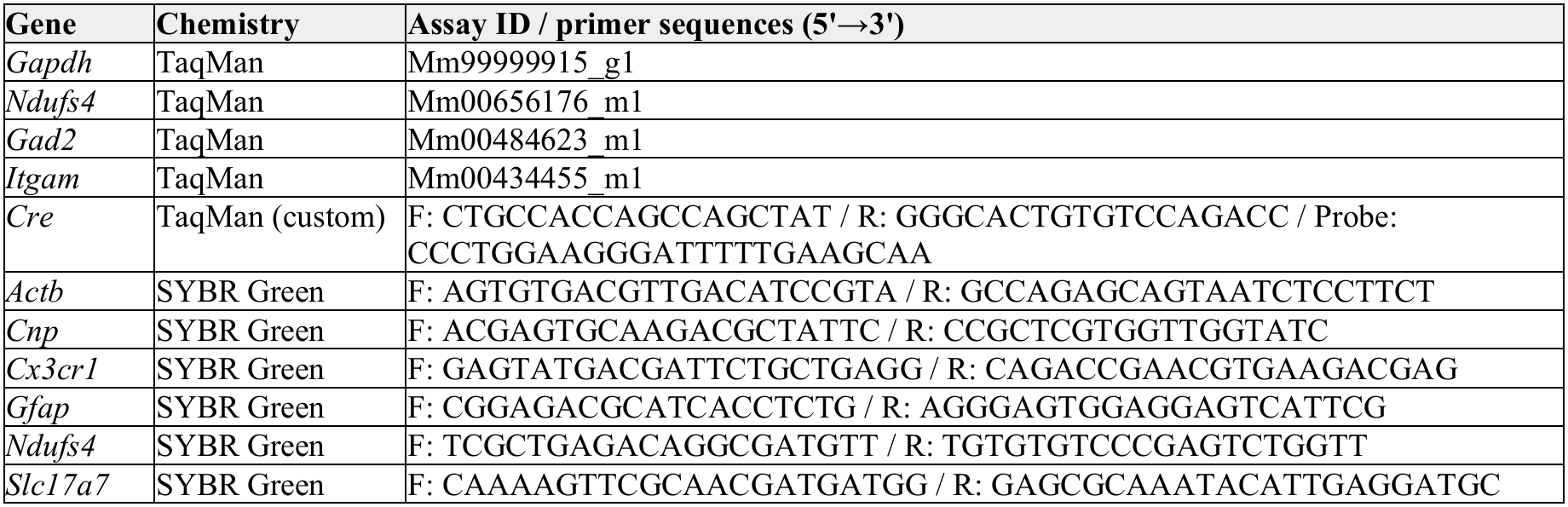
TaqMan assays and SYBR Green primers used for gene expression analysis.

## References

Aguilar, K., Comes, G., Canal, C., Quintana, A., Sanz, E., and Hidalgo, J. (2022). Microglial response promotes neurodegeneration in the Ndufs4 KO mouse model of Leigh syndrome. Glia 70, 2032–2044. 10.1002/glia.24234.

Bando, Y., Onuki, R., Katayama, T., Manabe, T., Kudo, T., Taira, K., and Tohyama, M. (2005). Double-strand RNA dependent protein kinase (PKR) is involved in the extrastriatal degeneration in Parkinson’s disease and Huntington’s disease. Neurochem Int 46, 11–18. 10.1016/j.neuint.2004.07.005.

Bolea, I., Gella, A., Sanz, E., Prada-Dacasa, P., Menardy, F., Bard, A.M., Machuca-Marquez, P., Eraso-Pichot, A., Modol-Caballero, G., Navarro, X., et al. (2019). Defined neuronal populations drive fatal phenotype in a mouse model of Leigh syndrome. Elife 8. 10.7554/eLife.47163.

Borgius, L., Restrepo, C.E., Leao, R.N., Saleh, N., and Kiehn, O. (2010). A transgenic mouse line for molecular genetic analysis of excitatory glutamatergic neurons. Molecular and cellular neurosciences 45, 245–257. 10.1016/j.mcn.2010.06.016.

Bose, A., Mouton-Liger, F., Paquet, C., Mazot, P., Vigny, M., Gray, F., and Hugon, J. (2011). Modulation of tau phosphorylation by the kinase PKR: implications in Alzheimer’s disease. Brain Pathol 21, 189–200. 10.1111/j.1750-3639.2010.00437.x.

Brooks, S.P., and Dunnett, S.B. (2009). Tests to assess motor phenotype in mice: a user’s guide. Nat Rev Neurosci 10, 519–529. 10.1038/nrn2652.

Bruchas, M.R., Schindler, A.G., Shankar, H., Messinger, D.I., Miyatake, M., Land, B.B., Lemos, J.C., Hagan, C.E., Neumaier, J.F., Quintana, A., et al. (2011). Selective p38alpha MAPK deletion in serotonergic neurons produces stress resilience in models of depression and addiction. Neuron 71, 498–511. 10.1016/j.neuron.2011.06.011.

Camandola, S., and Mattson, M.P. (2017). Brain metabolism in health, aging, and neurodegeneration. EMBO J 36, 1474–1492. 10.15252/embj.201695810.

Canal, C., Aguilar, K., Comes, G., Sanz, E., Giralt, M., Sanchis, P., and Hidalgo, J. (2026). Chronic central-targeted Interleukin-6 overexpression promotes hippocampal and cortical neuropathology in the Tg2576 mouse model of Alzheimer’s disease. Alzheimers Res Ther 18. 10.1186/s13195-026-02101-9.

Cheng, W.Y., Lee, X.Z., Lai, M.S., Ho, Y.S., and Chang, R.C. (2025). PKR modulates sterile systemic inflammation-triggered neuroinflammation and brain glucose metabolism disturbances. Front Immunol 16, 1469737. 10.3389/fimmu.2025.1469737.

Corra, S., Cerutti, R., Balmaceda, V., Viscomi, C., and Zeviani, M. (2022). Double administration of self-complementary AAV9NDUFS4 prevents Leigh disease in Ndufs4-/-mice. Brain 145, 3405–3414. 10.1093/brain/awac182.

Daneshgar, N., Leidinger, M.R., Le, S., Hefti, M., Prigione, A., and Dai, D.F. (2022). Activated microglia and neuroinflammation as a pathogenic mechanism in Leigh syndrome. Front Neurosci 16, 1068498. 10.3389/fnins.2022.1068498.

Dhir, A., Dhir, S., Borowski, L.S., Jimenez, L., Teitell, M., Rotig, A., Crow, Y.J., Rice, G.I., Duffy, D., Tamby, C., et al. (2018). Mitochondrial double-stranded RNA triggers antiviral signalling in humans. Nature 560, 238–242. 10.1038/s41586-018-0363-0.

Di Filippo, M., Tozzi, A., Arcangeli, S., de Iure, A., Durante, V., Di Gregorio, M., Gardoni, F., and Calabresi, P. (2016). Interferon-beta1a modulates glutamate neurotransmission in the CNS through CaMKII and GluN2A-containing NMDA receptors. Neuropharmacology 100, 98–105. 10.1016/j.neuropharm.2015.06.009.

Evans, H.T., Bodea, L.G., and Gotz, J. (2020). Cell-specific non-canonical amino acid labelling identifies changes in the de novo proteome during memory formation. Elife 9. 10.7554/eLife.52990.

Fadok, J.P., Dickerson, T.M., and Palmiter, R.D. (2009). Dopamine is necessary for cue-dependent fear conditioning. J Neurosci 29, 11089–11097. 10.1523/JNEUROSCI.1616-09.2009.

Fassone, E., and Rahman, S. (2012). Complex I deficiency: clinical features, biochemistry and molecular genetics. J Med Genet 49, 578–590. 10.1136/jmedgenet-2012-101159.

Fouche, B.R., Khumalo, S.G., Koopman, W.J.H., and Venter, M. (2026). Leigh Syndrome Pathomechanism Involves Region-Specific Innate Immune Activation in Ndufs4 Knockout Mice. Cell Mol Neurobiol 46, 42. 10.1007/s10571-026-01681-2.

Gal-Ben-Ari, S., Barrera, I., Ehrlich, M., and Rosenblum, K. (2018). PKR: A Kinase to Remember. Front Mol Neurosci 11, 480. 10.3389/fnmol.2018.00480.

Gella, A., Prada-Dacasa, P., Carrascal, M., Urpi, A., Gonzalez-Torres, M., Abian, J., Sanz, E., and Quintana, A. (2020). Mitochondrial Proteome of Affected Glutamatergic Neurons in a Mouse Model of Leigh Syndrome. Front Cell Dev Biol 8, 660. 10.3389/fcell.2020.00660.

Gorman, G.S., Chinnery, P.F., DiMauro, S., Hirano, M., Koga, Y., McFarland, R., Suomalainen, A., Thorburn, D.R., Zeviani, M., and Turnbull, D.M. (2016). Mitochondrial diseases. Nat Rev Dis Primers 2, 16080. 10.1038/nrdp.2016.80.

Gorman, G.S., Schaefer, A.M., Ng, Y., Gomez, N., Blakely, E.L., Alston, C.L., Feeney, C., Horvath, R., Yu-Wai-Man, P., Chinnery, P.F., et al. (2015). Prevalence of nuclear and mitochondrial DNA mutations related to adult mitochondrial disease. Ann Neurol 77, 753–759. 10.1002/ana.24362.

Hanaford, A., and Johnson, S.C. (2022). The immune system as a driver of mitochondrial disease pathogenesis: a review of evidence. Orphanet J Rare Dis 17, 335. 10.1186/s13023-022-02495-3.

Hanaford, A.R., Khanna, A., Truong, V., James, K., Chen, Y., Mulholland, M., Kayser, B., Liao, R.W., Sedensky, M., Morgan, P., et al. (2023). Peripheral macrophages drive CNS disease in the Ndufs4(-/-) model of Leigh syndrome. Brain Pathol 33, e13192. 10.1111/bpa.13192.

Hosseini, S., Michaelsen-Preusse, K., Grigoryan, G., Chhatbar, C., Kalinke, U., and Korte, M. (2020). Type I Interferon Receptor Signaling in Astrocytes Regulates Hippocampal Synaptic Plasticity and Cognitive Function of the Healthy CNS. Cell Rep 31, 107666. 10.1016/j.celrep.2020.107666.

Johnson, S.C., Yanos, M.E., Kayser, E.B., Quintana, A., Sangesland, M., Castanza, A., Uhde, L., Hui, J., Wall, V.Z., Gagnidze, A., et al. (2013). mTOR inhibition alleviates mitochondrial disease in a mouse model of Leigh syndrome. Science 342, 1524–1528. 10.1126/science.1244360.

Kang, R., and Tang, D. (2012). PKR-dependent inflammatory signals. Sci Signal 5, pe47. 10.1126/scisignal.2003511.

Keshavan, N., Mhaldien, L., Gilmour, K., and Rahman, S. (2024). Interferon Stimulated Gene Expression Is a Biomarker for Primary Mitochondrial Disease. Ann Neurol 96, 1185–1200. 10.1002/ana.27081.

Kim, Y., Park, J., Kim, S., Kim, M., Kang, M.G., Kwak, C., Kang, M., Kim, B., Rhee, H.W., and Kim, V.N. (2018). PKR Senses Nuclear and Mitochondrial Signals by Interacting with Endogenous Double-Stranded RNAs. Mol Cell 71, 1051–1063 e1056. 10.1016/j.molcel.2018.07.029.

Kruse, S.E., Watt, W.C., Marcinek, D.J., Kapur, R.P., Schenkman, K.A., and Palmiter, R.D. (2008). Mice with mitochondrial complex I deficiency develop a fatal encephalomyopathy. Cell Metab 7, 312–320. 10.1016/j.cmet.2008.02.004.

Lake, N.J., Compton, A.G., Rahman, S., and Thorburn, D.R. (2016). Leigh syndrome: One disorder, more than 75 monogenic causes. Ann Neurol 79, 190–203. 10.1002/ana.24551.

Lax, N.Z., Gorman, G.S., and Turnbull, D.M. (2017). Review: Central nervous system involvement in mitochondrial disease. Neuropathol Appl Neurobiol 43, 102–118. 10.1111/nan.12333.

Lee, H., Fenster, R.J., Pineda, S.S., Gibbs, W.S., Mohammadi, S., Davila-Velderrain, J., Garcia, F.J., Therrien, M., Novis, H.S., Gao, F., et al. (2020). Cell Type-Specific Transcriptomics Reveals that Mutant Huntingtin Leads to Mitochondrial RNA Release and Neuronal Innate Immune Activation. Neuron 107, 891–908 e898. 10.1016/j.neuron.2020.06.021.

Lee, J.J., Hsiao, Y.T., Huang, H.M., Ke, M.C., Tsao, S.T., Huang, Y.S., and Lin, T.K. (2026). Inhibition of STING protects the neuroretina in an Ndufs4 knockout mouse model of mitochondrial complex 1 deficiency. Exp Eye Res 273, 111238. 10.1016/j.exer.2026.111238.

Lei, Y., Guerra Martinez, C., Torres-Odio, S., Bell, S.L., Birdwell, C.E., Bryant, J.D., Tong, C.W., Watson, R.O., West, L.C., and West, A.P. (2021). Elevated type I interferon responses potentiate metabolic dysfunction, inflammation, and accelerated aging in mtDNA mutator mice. Sci Adv 7. 10.1126/sciadv.abe7548.

Lepelley, A., Della Mina, E., Van Nieuwenhove, E., Waumans, L., Fraitag, S., Rice, G.I., Dhir, A., Fremond, M.L., Rodero, M.P., Seabra, L., et al. (2021). Enhanced cGAS-STING-dependent interferon signaling associated with mutations in ATAD3A. J Exp Med 218. 10.1084/jem.20201560.

Liu, L., Zhang, K., Sandoval, H., Yamamoto, S., Jaiswal, M., Sanz, E., Li, Z., Hui, J., Graham, B.H., Quintana, A., and Bellen, H.J. (2015). Glial lipid droplets and ROS induced by mitochondrial defects promote neurodegeneration. Cell 160, 177–190. 10.1016/j.cell.2014.12.019.

Lopez-Atalaya, J.P., and Bhojwani-Cabrera, A.M. (2025). Type I interferon signalling and interferon-responsive microglia in health and disease. FEBS J 292, 5921–5940. 10.1111/febs.70126.

Lopez-Grancha, M., Bernardelli, P., Moindrot, N., Genet, E., Vincent, C., Roudieres, V., Krick, A., Sabuco, J.F., Machnik, D., Ibghi, D., et al. (2021). A Novel Selective PKR Inhibitor Restores Cognitive Deficits and Neurodegeneration in Alzheimer Disease Experimental Models. J Pharmacol Exp Ther 378, 262–275. 10.1124/jpet.121.000590.

Love, M.I., Huber, W., and Anders, S. (2014). Moderated estimation of fold change and dispersion for RNA-seq data with DESeq2. Genome Biol 15, 550. 10.1186/s13059-014-0550-8.

Luna, M., González-Torres, M., Girona, M., and Quintana, A. (2025). Mitochondrial Nucleic Acids as Innate Immune Activators: Molecular Mechanisms and Pathophysiological Impact. In The Oxford Handbook of Mitochondria, E.A. Jonas, ed. (Oxford University Press). 10.1093/oxfordhb/9780190932183.013.27.

Magro, G., Laterza, V., and Tosto, F. (2025). Leigh Syndrome: A Comprehensive Review of the Disease and Present and Future Treatments. Biomedicines 13. 10.3390/biomedicines13030733.

Marques, E., Kramer, R., and Ryan, D.G. (2024). Multifaceted mitochondria in innate immunity. NPJ Metab Health Dis 2, 6. 10.1038/s44324-024-00008-3.

McElroy, G.S., Reczek, C.R., Reyfman, P.A., Mithal, D.S., Horbinski, C.M., and Chandel, N.S. (2020). NAD+ Regeneration Rescues Lifespan, but Not Ataxia, in a Mouse Model of Brain Mitochondrial Complex I Dysfunction. Cell Metab 32, 301–308 e306. 10.1016/j.cmet.2020.06.003.

McFarland, R., Taylor, R.W., and Turnbull, D.M. (2010). A neurological perspective on mitochondrial disease. Lancet Neurol 9, 829–840. 10.1016/S1474-4422(10)70116-2.

Miura, Y., Li, M.Y., Birey, F., Ikeda, K., Revah, O., Thete, M.V., Park, J.Y., Puno, A., Lee, S.H., Porteus, M.H., and Pasca, S.P. (2020). Generation of human striatal organoids and cortico-striatal assembloids from human pluripotent stem cells. Nat Biotechnol 38, 1421–1430. 10.1038/s41587-020-00763-w.

Muller, U., Steinhoff, U., Reis, L.F., Hemmi, S., Pavlovic, J., Zinkernagel, R.M., and Aguet, M. (1994). Functional role of type I and type II interferons in antiviral defense. Science 264, 1918–1921. 10.1126/science.8009221.

Nazmi, A., Field, R.H., Griffin, E.W., Haugh, O., Hennessy, E., Cox, D., Reis, R., Tortorelli, L., Murray, C.L., Lopez-Rodriguez, A.B., et al. (2019). Chronic neurodegeneration induces type I interferon synthesis via STING, shaping microglial phenotype and accelerating disease progression. Glia 67, 1254–1276. 10.1002/glia.23592.

Okuda, S., Watanabe, Y., Moriya, Y., Kawano, S., Yamamoto, T., Matsumoto, M., Takami, T., Kobayashi, D., Araki, N., Yoshizawa, A.C., et al. (2017). jPOSTrepo: an international standard data repository for proteomes. Nucleic Acids Res 45, D1107–D1111. 10.1093/nar/gkw1080.

Onuki, R., Bando, Y., Suyama, E., Katayama, T., Kawasaki, H., Baba, T., Tohyama, M., and Taira, K. (2004). An RNA-dependent protein kinase is involved in tunicamycin-induced apoptosis and Alzheimer’s disease. EMBO J 23, 959–968. 10.1038/sj.emboj.7600049.

Perry, E.A., Bennett, C.F., Luo, C., Balsa, E., Jedrychowski, M., O’Malley, K.E., Latorre-Muro, P., Ladley, R.P., Reda, K., Wright, P.M., et al. (2021). Tetracyclines promote survival and fitness in mitochondrial disease models. Nat Metab 3, 33–42. 10.1038/s42255-020-00334-y.

Prada-Dacasa, P., Urpi, A., Sanchez-Benito, L., Bianchi, P., and Quintana, A. (2020). Measuring Breathing Patterns in Mice Using Whole-body Plethysmography. Bio Protoc 10, e3741. 10.21769/BioProtoc.3741.

Puighermanal, E., Luna-Sanchez, M., Gella, A., van der Walt, G., Urpi, A., Royo, M., Tena-Morraja, P., Appiah, I., de Donato, M.H., Menardy, F., et al. (2024). Cannabidiol ameliorates mitochondrial disease via PPARgamma activation in preclinical models. Nat Commun 15, 7730. 10.1038/s41467-024-51884-8.

Quintana, A., Kruse, S.E., Kapur, R.P., Sanz, E., and Palmiter, R.D. (2010). Complex I deficiency due to loss of Ndufs4 in the brain results in progressive encephalopathy resembling Leigh syndrome. Proc Natl Acad Sci U S A 107, 10996–11001. 10.1073/pnas.1006214107.

Quintana, A., Zanella, S., Koch, H., Kruse, S.E., Lee, D., Ramirez, J.M., and Palmiter, R.D. (2012). Fatal breathing dysfunction in a mouse model of Leigh syndrome. J Clin Invest 122, 2359–2368. 10.1172/JCI62923.

Rahman, J., and Rahman, S. (2018). Mitochondrial medicine in the omics era. Lancet 391, 2560–2574. 10.1016/S0140-6736(18)30727-X.

Roy, E.R., Chiu, G., Li, S., Propson, N.E., Kanchi, R., Wang, B., Coarfa, C., Zheng, H., and Cao, W. (2022). Concerted type I interferon signaling in microglia and neural cells promotes memory impairment associated with amyloid beta plaques. Immunity 55, 879–894 e876. 10.1016/j.immuni.2022.03.018.

Roy, E.R., Wang, B., Wan, Y.W., Chiu, G., Cole, A., Yin, Z., Propson, N.E., Xu, Y., Jankowsky, J.L., Liu, Z., et al. (2020). Type I interferon response drives neuroinflammation and synapse loss in Alzheimer disease. J Clin Invest 130, 1912–1930. 10.1172/JCI133737.

Sanchez-Benito, L., Gonzalez-Torres, M., Fernandez-Gonzalez, I., Cutando, L., Royo, M., Compte, J., Vila, M., Jurado, S., Sanz, E., and Quintana, A. (2025). Dysfunctional LHX6 pallido-subthalamic projections mediate epileptic events in a mouse model of Leigh syndrome. J Clin Invest 135. 10.1172/JCI187571.

Sanz, E., Bean, J.C., Carey, D.P., Quintana, A., and McKnight, G.S. (2019). RiboTag: Ribosomal Tagging Strategy to Analyze Cell-Type-Specific mRNA Expression In Vivo. Curr Protoc Neurosci 88, e77. 10.1002/cpns.77.

Sanz, E., Yang, L., Su, T., Morris, D.R., McKnight, G.S., and Amieux, P.S. (2009). Cell-type-specific isolation of ribosome-associated mRNA from complex tissues. Proc Natl Acad Sci U S A 106, 13939–13944. 10.1073/pnas.0907143106.

Sauer, J.D., Sotelo-Troha, K., von Moltke, J., Monroe, K.M., Rae, C.S., Brubaker, S.W., Hyodo, M., Hayakawa, Y., Woodward, J.J., Portnoy, D.A., and Vance, R.E. (2011). The N-ethyl-N-nitrosourea-induced Goldenticket mouse mutant reveals an essential function of Sting in the in vivo interferon response to Listeria monocytogenes and cyclic dinucleotides. Infect Immun 79, 688–694. 10.1128/IAI.00999-10.

Seibenhener, M.L., and Wooten, M.C. (2015). Use of the Open Field Maze to measure locomotor and anxiety-like behavior in mice. J Vis Exp, e52434. 10.3791/52434.

Stokes, J.C., Bornstein, R.L., James, K., Park, K.Y., Spencer, K.A., Vo, K., Snell, J.C., Johnson, B.M., Morgan, P.G., Sedensky, M.M., et al. (2022). Leukocytes mediate disease pathogenesis in the Ndufs4(KO) mouse model of Leigh syndrome. JCI Insight 7. 10.1172/jci.insight.156522.

Suomalainen, A., and Battersby, B.J. (2018). Mitochondrial diseases: the contribution of organelle stress responses to pathology. Nat Rev Mol Cell Biol 19, 77–92. 10.1038/nrm.2017.66.

Taniguchi, H., He, M., Wu, P., Kim, S., Paik, R., Sugino, K., Kvitsiani, D., Fu, Y., Lu, J., Lin, Y., et al. (2011). A resource of Cre driver lines for genetic targeting of GABAergic neurons in cerebral cortex. Neuron 71, 995–1013. 10.1016/j.neuron.2011.07.026.

Tible, M., Mouton Liger, F., Schmitt, J., Giralt, A., Farid, K., Thomasseau, S., Gourmaud, S., Paquet, C., Rondi Reig, L., Meurs, E., et al. (2019). PKR knockout in the 5xFAD model of Alzheimer’s disease reveals beneficial effects on spatial memory and brain lesions. Aging Cell 18, e12887. 10.1111/acel.12887.

Unwin, R.D., Griffiths, J.R., and Whetton, A.D. (2010). Simultaneous analysis of relative protein expression levels across multiple samples using iTRAQ isobaric tags with 2D nano LC-MS/MS. Nat Protoc 5, 1574–1582. 10.1038/nprot.2010.123.

van de Wal, M.A.E., Adjobo-Hermans, M.J.W., Keijer, J., Schirris, T.J.J., Homberg, J.R., Wieckowski, M.R., Grefte, S., van Schothorst, E.M., van Karnebeek, C., Quintana, A., and Koopman, W.J.H. (2022). Ndufs4 knockout mouse models of Leigh syndrome: pathophysiology and intervention. Brain 145, 45–63. 10.1093/brain/awab426.

Wang, R., Yang, B., and Zhang, D. (2011). Activation of interferon signaling pathways in spinal cord astrocytes from an ALS mouse model. Glia 59, 946–958. 10.1002/glia.21167.

Warren, E.B., Gordon-Lipkin, E.M., Cheung, F., Chen, J., Mukherjee, A., Apps, R., Tsang, J.S., Jetmore, J., Schlein, M.L., Kruk, S., et al. (2023). Inflammatory and interferon gene expression signatures in patients with mitochondrial disease. J Transl Med 21, 331. 10.1186/s12967-023-04180-w.

West, A.P., Khoury-Hanold, W., Staron, M., Tal, M.C., Pineda, C.M., Lang, S.M., Bestwick, M., Duguay, B.A., Raimundo, N., MacDuff, D.A., et al. (2015). Mitochondrial DNA stress primes the antiviral innate immune response. Nature 520, 553–557. 10.1038/nature14156.

Yang, Y.L., Reis, L.F., Pavlovic, J., Aguzzi, A., Schafer, R., Kumar, A., Williams, B.R., Aguet, M., and Weissmann, C. (1995). Deficient signaling in mice devoid of double-stranded RNA-dependent protein kinase. EMBO J 14, 6095–6106. 10.1002/j.1460-2075.1995.tb00300.x.

Zink, A., Dai, D.F., Wittich, A., Henke, M.T., Pedrotti, G., Heiduschka, S., Santamaria, G., Pentimalli, T.M., Brueser, C., Notopoulou, S., et al. (2026). Pluripotent stem-cell-based screening uncovers sildenafil as a mitochondrial disease therapy. Cell 189, 1656–1679 e1642. 10.1016/j.cell.2026.02.008.

